# Live-cell transcriptomics with engineered virus-like particles

**DOI:** 10.1101/2024.10.01.616098

**Authors:** Mohamad Ali Najia, Jacob Borrajo, Anna Le, FuNien Tsai, Jeremy Y. Huang, Linda G. Griffith, George Q. Daley, Paul C. Blainey

## Abstract

The transcriptional state of a cell provides a multi-parameter representation of gene expression programs that reflect its identity and phenotype. However, current transcriptomic profiling technologies result in destruction of the biological sample, preventing direct analysis of transcriptional dynamics in the same living cells over time. Here, we developed a synthetic RNA export system called cellular ‘self-reporting’ to address this fundamental technological limitation. We repurposed the murine leukemia virus retroviral protein Gag to enable diverse types of immortalized and primary mammalian cells to package cellular RNA molecules in virus-like particles (VLPs) for export into the extracellular environment. We applied self-reporting to interrogate the transcriptome-wide dynamics that occur during neuronal differentiation from induced pluripotent stem cells and detected gene expression profiles from individual live cells. Leveraging this genetically encodable approach, we expanded the capabilities of self-reporting through molecular engineering of VLP components. Pseudotyping VLPs with epitope-tagged envelope proteins enabled multiplexed selective live-cell readout of transcriptional states from heterogeneous co-cultures. Furthermore, structure-guided protein engineering of Gag fusions with human RNA binding domains improved the mRNA representation in self-reporting readouts and enabled the directed export of libraries of synthetic barcode transcripts. Taken together, this work establishes self-reporting as a facile and broadly enabling technology for live-cell, transcriptome-scale profiling of dynamic processes across diverse cell types and biological applications.

## INTRODUCTION

The dynamic regulation of gene expression is integral to cellular identity and function. As such, the compendium of expressed genes in a cell is an important source of molecular information relevant to cellular processes occurring during both normal development and pathogenesis. Over the last 15 years, unbiased gene expression profiling, particularly via high-throughput RNA-sequencing (RNA-seq), has become a cornerstone of modern biomedical science^1–3^. Research interest in the dynamic evolution of transcriptional states over the course of diverse biological processes has also been made clear by the proliferation of computational “pseudotime” methods inferring underlying temporal evolution^4–10^. However, current state-of-the-art approaches for RNA profiling, such as single-cell RNA-seq, almost exclusively result in destruction of cells in the biological sample, as cells are lysed for RNA extraction or transcripts are retrieved from a few cells using invasive micromechanical procedures at each time point^11,12^. The destructive nature of current RNA-seq technologies prevents the integration of multiple functional readouts on the same biological sample, hindering the ability to study dynamic cellular phenomena.

Indeed, targeted approaches to study time-dependent gene expression have been an active area of technology development. The advent of genetically encodable fluorescent proteins provided an early framework for enabling true time series analysis of individual gene products using optical microscopy. Several groups have further pioneered optical methods that use endogenous or exogenous probes to monitor a few native or tagged target transcripts at a time^13–23^. TIVA enables transcriptome-wide RNA capture in living cells but has not yet been reported for longitudinal profiling^24^. More recently, other molecular recording methods have allowed information about the age of transcripts or prior transcriptional states to be encoded in the DNA or RNA of living systems for readout at a single end point^25–28^. While useful, these methods require a prior transcriptional hypothesis, have limited multiplexing capacity and dynamic range, and do not retain the living biological sample after the measurement. Overall, the limitations of these approaches further highlight the need for broadly accessible technologies that allow true live-cell, longitudinal monitoring of transcriptome-wide gene expression to study dynamic cellular phenomena.

Inspired by natural systems, we sought to overcome the limitations of conventional RNA-seq by leveraging synthetic biology and protein engineering to create an RNA export pathway that enables mammalian cells to “self-report” their transcriptional states in real time. Exosomes are membrane-bound extracellular vesicles that may contain cellular RNAs and are naturally released from mammalian cells. However, exosome biogenesis and function remain largely enigmatic and are highly cell-type and context dependent^29^. Instead, we focused on retroviruses, which evolved to package and export their RNA genomes from mammalian cells in virions. The murine leukemia virus (MLV), in particular, is among the simplest and most well-characterized retroviruses, whose genome comprises three genes: *gag*, *pol*, and *env*. Expression of Gag, the core structural polyprotein, has long been known to be sufficient for the formation of replication-incompetent, virus-like particles (VLPs) in mammalian hosts^30^. The ability to produce VLPs has ushered a broad range of technological applications, including vaccines^31^, measurement of protein-protein interactions^32^, and cellular export and delivery of proteins or synthetic nucleic acids^33–36^. While many applications of VLPs have leveraged the ability to augment the payload that is packaged, RNA remains an essential structural component of a retrovirus particle^37,38^. However, in the absence of a viral genome or other *cis-*acting, viral packaging signals, Gag assembly is not impaired and the resulting particles still contain RNA^36,39,40^. These findings and microarray-based profiling of MLV Gag VLPs suggested that cellular RNAs can be incorporated in lieu of the viral genomic RNA^41^. We reasoned that retroviral machinery is an attractive substrate for a genetically encoded synthetic system because it is orthogonal to mammalian host biology, allowing for precise molecular engineering to tailor the properties of VLPs and control their activation in time and space. Thus, we hypothesized that MLV Gag could be repurposed as a useful tool for unbiased export of cellular RNAs in VLPs to the extracellular environment, enabling non-destructive, transcriptome-wide profiling of RNAs collected at a distance from the self-reporting cell (**Figure 1A-B**). We envision that live-cell transcriptional profiling will further our understanding of dynamic biological phenomena from transcriptome-scale measurements.

**Figure 1.**
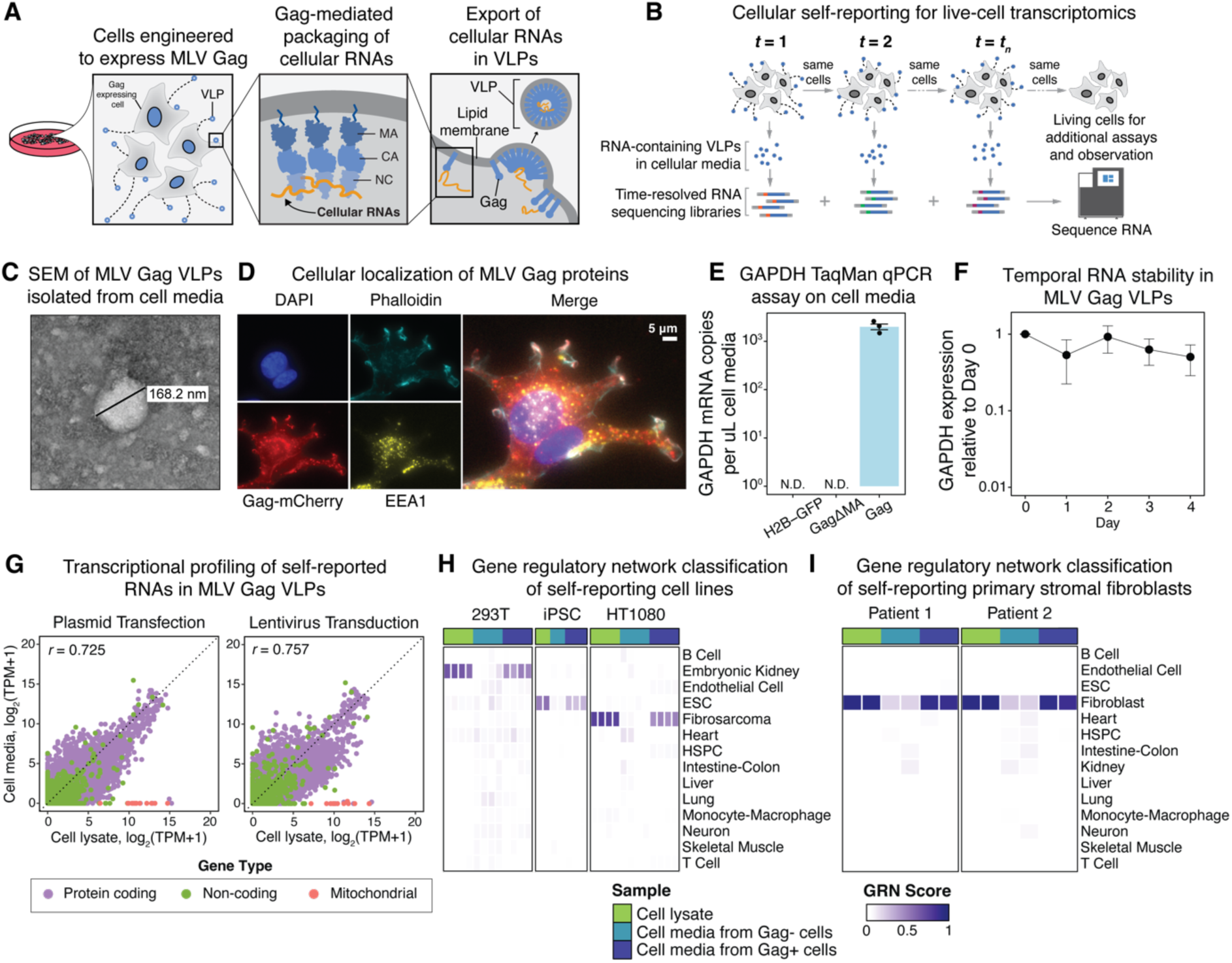
MLV Gag enables live-cell transcriptomics. **(A)** Schematic of MLV Gag-mediated export of cellular RNAs in VLPs. **(B)** Experimental schematic outlining that engineered cells self-report their transcriptional states within VLPs to enable live-cell, temporal transcriptome-wide measurements. **(C)** Scanning electron microscopy of purified VLPs from cellular media. **(D)** Immunofluorescence microscopy of HEK293T cells expressing Gag-mCherry fusion protein to track subcellular localization of VLP production. Cells were co-stained for actin filaments with phalloidin, nuclei with DAPI, and endosomal marker EEA1. Scale bar corresponds to 5 µm. **(E)** TaqMan qPCR assay to quantify GAPDH expression from purified cellular media of HEK293T cells transfected with either H2B-GFP, GagΔMA or Gag expression constructs. Each point represents n=3 transfection independent replicates and error bars represent standard deviation. **(F)** TaqMan qPCR assay to quantify GAPDH from VLPs incubated at 37°C in cellular media over four days. **(G)** Scatterplot of transcript abundance detected from cellular lysate and VLPs purified from cell media. Each point represents a gene detected from RNA-seq (colored by the transcript type: protein coding, non-coding, or mitochondrial) and the values are averaged over n=3 matched replicates from cellular lysate and VLPs. HEK293T cells were engineered to self-report cellular RNAs in VLPs via transient plasmid transfection (left) or lentiviral transduction (right) of MLV *gag*. TPM = transcripts per million. **(H-I)** CellNet classification heatmap of cellular lysate and VLP-derived RNA from lentivirus infected, constitutively self-reporting HEK293T cells, HT1080 cells, iPS cells **(H)** and primary patient fibroblasts **(I)**. The gene regulatory network (GRN) score represents the degree to which the sample recapitulates the GRN of the known cell types within the CellNet classifier.

## RESULTS

### MLV Gag exports cellular RNAs stabilized in VLPs

To evaluate the utility of retroviral machinery as the basis of a synthetic RNA export pathway, we focused on MLV Gag since it is the minimal essential gene required to form VLPs. We synthesized the full coding sequence consisting of the N-terminal matrix (MA, p15) domain, the central capsid (CA, p30) domain and C-terminal nucleocapsid (NC, p10) domain (**Supplemental Table 1**), and transfected it into HEK293T cells. Since Gag localization and assembly on lipid membranes is mediated through the MA domain, we generated a Gag truncation mutant lacking the MA domain (GagΔMA) as a negative control for VLP formation^42,43^. We detected MLV Gag p30 protein in the culture medium of cells transfected with full-length *gag*, but not in the medium of cells expressing GagΔMA (**Figure S1A**), as expected. Furthermore, scanning electron microscopy revealed the formation of VLPs approximately 160 nm in diameter in the culture media of cells expressing Gag (**Figure 1C**). We also observed that Gag C-terminally fused to mCherry formed distinct punctate foci within cells that co-localized with the endosomal marker EEA1 (**Figure 1D**). Consistent with prior reports, our data implicate endosomal pathways in VLP egress^30,44,45^. The utilization of this conserved host machinery suggests that VLPs could be produced in diverse types of mammalian cells.

We then asked whether cellular RNAs are packaged and exported in VLPs. We designed a TaqMan-based qPCR assay to quantitatively measure the abundance of a ubiquitously expressed gene, *GAPDH.* Purification of cell media from HEK293T cells transfected with full-length *gag* revealed detectable *GAPDH* transcripts, yet *GAPDH* was not detected in the media of cells transfected with negative control expression vectors encoding GagΔMA or H2B-GFP (**Figure 1E**). To determine the breadth of cellular RNAs associated with VLPs and confirm these RNAs were indeed packaged inside the VLPs, we treated the cell media with benzonase endonuclease, which non-specifically degrades RNA and DNA, and performed RNA-seq^46^ on the treated samples (**Figure S1B)**. We observed a 10-fold enrichment of RNA-seq reads that aligned across the human transcriptome from Gag-expressing cells compared to GagΔMA-expressing control cells (**Figure S1C**), demonstrating that VLPs can protect a broad range of cellular transcripts from nuclease activity outside the cellular environment. In another test of RNA stabilization by VLPs, we observed that *GAPDH* transcripts were reliably detected in cell media isolated from Gag-expressing cells and incubated at 37°C over 4 days (**Figure 1F**). These results collectively underscore that MLV Gag-based VLPs can stabilize their RNA payload for analysis at a later time.

To further characterize the RNAs exported within VLPs, we performed full-length RNA-seq on VLPs harvested from the culture media of HEK293T cells transfected with *gag*. We detected on average 5,171 unique genes from purified VLP samples, compared to on average 7,754 genes from coupled cell lysate samples (**Figure S2A**), underscoring the high-dimensional transcriptional information that can be derived from self-reported RNA (**Figure S2B**). Importantly, the abundance of individual RNA transcripts detected from VLPs was generally concordant to traditional RNA-seq measurements on cell lysate (Pearson *r =* 0.725, **Figure 1G**), suggesting that transcriptional profiling of VLP-derived RNA could be quantitatively reflective of the cellular transcriptome. We noted, however, that while protein-coding and non-coding transcripts were represented in VLPs, all 13 known mitochondrial genes were absent (**Figure 1G**). This observation is consistent with the physical isolation of mitochondrial transcripts from the cytosol where Gag assembly occurs. Replicate reproducibility of VLP-derived RNAs was also highly robust and comparable to that of cell lysate-based RNA-seq (Pearson *r* = 0.897 for VLPs and *r* = 0.911 for cell lysate, **Figure S2C-D**). These data collectively establish that MLV Gag can enable living mammalian cells to “self-report” cellular transcripts via VLPs for live-cell transcriptomics.

### Diverse cell types can constitutively self-report their transcriptional state via VLPs

While our proof-of-principle data were based on transient transfection of *gag*, we next sought to engineer mammalian cells to constitutively express Gag in order to allow longitudinal live-cell studies of transcriptional dynamics. Lentiviral transduction of *gag* into HEK293T cells enabled secretion of VLPs into the cell media (**Figure S3A**). The relative transcript abundances detected in the media of cells constitutively expressing Gag also correlated with transcript abundances in cell lysates (Pearson *r* = 0.757), indicating that Gag expression from single genomic integrants results in comparable performance as plasmid transfection (**Figure 1G**). We generalized these findings beyond HEK293T cells by showing that HT1080 fibrosarcoma cells were also able to constitutively produce VLPs following lentiviral transfer of *gag* (**Figure S3A**). Extending these findings further, we then asked whether stem cell populations, such as induced pluripotent stem (iPS) cells, were also amenable for constitutive VLP production. Benzonase treatment of the cell media from Gag-expressing iPS cells demonstrated the production of nuclease-protected RNAs in the media that strongly reflected embryonic stem cell gene regulatory networks (GRNs), but the equivalent was not observed from control GagΔMA-expressing iPS cells (**Figure S3B**). In fact, we found that all constitutive Gag-expressing cell lines tested exhibited substantial levels of transcriptome export, and that transcript abundances from VLP-derived RNAs were highly concordant with transcript abundances from cell lysate (**Figure 1G, Figure S3C**). Importantly, we observed that constitutive expression of Gag was minimally perturbative across each cell line, evidenced by normal morphology, growth rates, and differential gene expression analysis of the cellular transcriptome, in which the transgenes were unsurprisingly found to be the most prominent differential signals (**Figure S3D-F**).

We then assessed whether the VLP-packaged RNA molecules from Gag-expressing cell lines were sufficiently reflective of the host transcriptome in order to distinguish and classify different cell types. To do so, we performed RNA-seq on VLP-derived RNAs obtained from each cell line over 48 hours then applied CellNet^47^ to ascertain active GRNs and predict the biological identity of each sample. The self-reported transcriptomes supported specific classification to the expected cellular identities, mirroring the classification performance of conventional cell lysate RNA-seq controls (**Figure 1H**). Finally, we observed similar GRN classification performance from Gag-expressing primary stromal fibroblasts across multiple donors, supporting that Gag-based self-reporting of transcriptional states in VLPs is applicable beyond immortalized cell lines (**Figure 1I**). These data demonstrate that VLP-derived RNAs enable a high-dimensional transcriptional readout amenable for gene regulatory network analysis and reflect the cellular transcriptome across diverse primary and immortalized cell types.

### Self-reported transcriptional signals are detectable across a broad range of timescales and in individual cells

We then leveraged our constitutive Gag-expressing cells to characterize the dynamics of RNA export in VLPs. First, we assessed self-reported transcriptional profiles as a function of the amount of time allowed for VLP secretion into the cell media (**Figure S4A**). We were able to detect VLP-derived RNA transcripts significantly above background in as little as three hours of VLP production (**Figure S4B-D**). CellNet classification of self-reported RNAs collected over various sampling windows further corroborated that the exported transcripts reflected cellular gene expression (**Figure S4B**). The classification performance of VLP-derived RNAs improved with longer durations of VLP production, generally plateauing at 48 hours (**Figure S4C**) and was associated with the number of genes detected per sampling interval (**Figure S4D**).

The sensitivity demonstrated in these experiments prompted us to characterize RNA export from individual cells. We engineered HEK293T cells with a stably-integrated, doxycycline (dox)-inducible Gag expression vector and FACS-isolated single cells in independent wells (**Figure S5A**). After culturing these cells for 24 hours, we prepared RNA-seq libraries from the single cells and associated cell media. We observed more cellular transcripts from cells with induced Gag expression compared to wild-type or uninduced cells, indicating that self-reported RNA profiles can be detected from single cells (**Figure S5B**). To further improve signal detection from cultures with few cells, we engineered HT1080 cells to constitutively express Gag and assessed self-reported RNAs from single cells or 100-cell cultures grown over 24 hours. We measured 1,127.0±830.5 transcripts in the media of single cells, compared to 10,301.9±1145.5 transcripts in the media of 100 cells (**Figure S5C**). Notably, the transcript abundances of genes detected in the media of single HT1080 cells correlated with the corresponding single-cell lysate measurements (**Figure S5D**) as well as measurements from 100 cells (**Figure S5E**), consistent with our data on bulk populations. These findings highlight our ability to detect exported RNA from single cells, and overall demonstrate the breadth of temporal and cell culture scales that are enabled by self-reporting.

### Live-cell transcriptional dynamics of neurogenesis modeled from pluripotent stem cells

Since we established that sampling VLPs from the media of Gag-expressing cells at a single time point reflects the cellular transcriptome, we next sought to investigate whether longitudinal sampling of VLPs could capture transcriptional dynamics within living cells over time (**Figure 1B**). iPS cells serve as useful *in vitro* models to study human development. We therefore utilized our self-reporting iPS cells to interrogate transcriptional changes over the course of cellular differentiation processes. Specifically, we focused on neurogenesis given that this process is fundamental to human brain development and neurological disorders. Furthermore, en route to becoming post-mitotic neurons, these differentiating cells progressively attenuate proliferation (**Figure S6A**) and thus cannot be subsampled for destructive timepoint assays indefinitely. The sampling constraint of non-cycling cell populations, including neurons, underscores the value of non-destructive, live-cell assays. We utilized iNGN iPS cells^48^, which leverage dox-inducible control of Neurogenin-1 and Neurogenin-2 transcription factors to facilitate the development of neurons with bipolar morphology within four days (**Figure S6B-C**). We introduced Gag into iNGN iPS cells via piggyBac transposition and verified that the cells can self-report their transcriptional state within VLPs (**Figure S6D-E**), consistent with our prior data in a separate iPS cell line (**Figure 1H**). We further validated that constitutive Gag expression did not negatively impede the differentiation process, evidenced by uniform bipolar morphology and surface expression of neural marker NCAM1 compared to GagΔMA-expressing cells (**Figure S6F-G**).

To measure transcriptional dynamics during neurogenesis, we induced the expression of Neurogenin-1 and Neurogenin-2 in iNGN iPSCs and differentiated these cells into bipolar-like neurons over the course of four days (**Figure 2A**). We collected samples for RNA-seq before induction and every 24 hours of the differentiation by repeatedly collecting media and purifying VLPs produced from the same cell population. To enable comparisons between self-reporting and traditional RNA-seq, we prepared additional cultures for destructive lysis at each time point.

**Figure 2.**
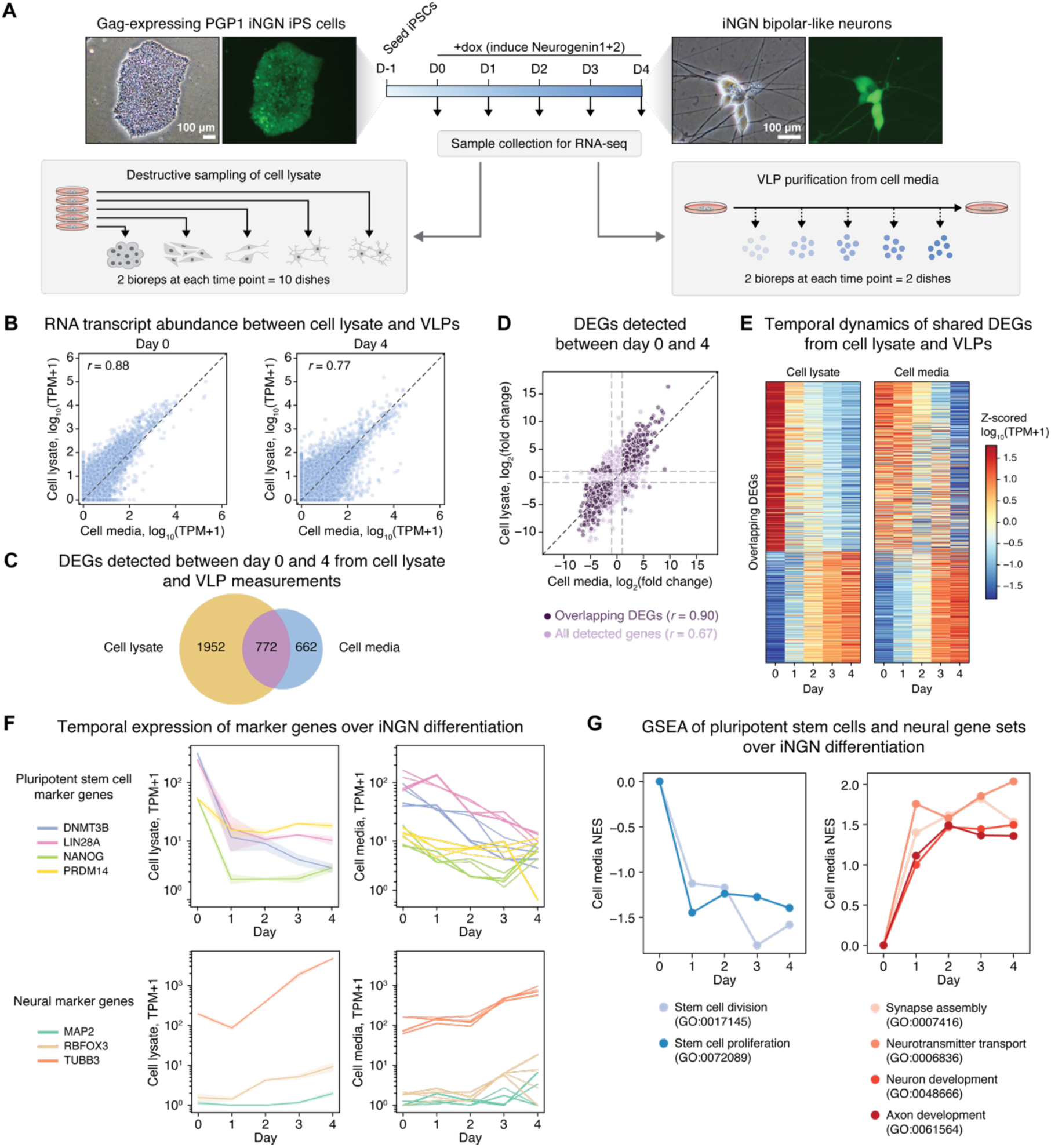
Live-cell transcriptional dynamics of neurogenesis. **(A)** Experimental workflow to longitudinally profile transcriptional dynamics during neuronal differentiation via self-reporting of cellular RNAs in VLPs. PGP1 iNGN iPS cells constitutively expressing Gag-2A-mNeon were differentiated into neurons by doxycycline induction of Neurogenin-1 and Neurogenin-2 over 4 days. Samples for RNA-seq were collected either destructively from cell lysate or non-destructively from culture media. Brightfield and mNeon (green) images show cells at days 0 and 4 of differentiation. **(B)** Gene expression concordance between cell lysates and VLPs from cell media at days 0 (left) and 4 (right) of neuronal differentiation. Log-transformed transcripts per million (TPM) were averaged across replicates (n=2 for cellular lysates, n=4 for cellular media). Pearson correlation coefficients *r* are presented in each plot. **(C)** Overlap in differentially expressed genes (DEGs) detected in cellular lysate and VLP samples. DEGs were defined as genes with log2(fold change) > 1 or log2(fold change) < −1 and false discovery rate (FDR)-adjusted *P* value below 0.01. **(D)** Scatterplot of fold changes in gene expression between day 0 and day 4 for all detected genes (light purple) and DEGs detected in both cellular lysates and corresponding VLPs (dark purple). Pearson correlation coefficients *r* are presented in each plot. **(E)** Heatmap of z-scored log-transformed TPMs. Genes are ordered by decreasing log2 fold change in cellular lysate. **(F)** Gene expression of stem cell (top) and neural (bottom) markers across 4-day differentiation in cellular lysate (top, mean ± SEM of n=2 replicates) and VLPs (bottom, each line showing one of n=4 replicates). **(G)** Line plots of normalized enrichment scores (NES) for select gene ontology (GO) terms describing stem cell states (left) and neuron development (right) across 4 days of differentiation. Gene set enrichment analysis (GSEA) was performed on ranked log2 fold changes in gene expression between each time point to day 0 from VLPs.

We first assessed the technical performance of cellular self-reporting in differentiating iPSCs. We found that gene expression profiles obtained from cell media were highly correlated with measurements from cell lysates and highly reproducible among replicates throughout the four-day time course (**Figure 2B, S7B-C**). Furthermore, principal component analysis (PCA) revealed distinct gene expression profiles across time for both cell lysate and cell media samples, suggesting that self-reporting can detect transcriptional changes occurring during differentiation (**Figure S7D**).

We next performed differential gene expression analysis to identify temporally dynamic genes during iNGN differentiation. We observed that 40% of the differentially expressed genes detected in cell lysate samples between day 0 and day 4 were also detected in cell media from live cultures (**Figure 2C, Supplemental Table 2A-B**). Interestingly, the gene expression fold changes from VLP-derived RNAs were highly concordant with measurements in cell lysate (**Figure 2D**), and in fact, can be used to predict differentially expressed genes in lysate with moderate accuracy (ROC AUC = 0.62) (**Figure S7E**). Among the shared differentially expressed genes, we also observed similar temporal patterns of gene expression across all four days of differentiation in both cell lysate and VLP-derived RNA samples (**Figure 2E**). We then specifically compared the expression levels of stem cell and neuronal marker genes in cell lysate and media samples over the course of neurogenesis. As expected, we detected time-dependent attenuation in the expression of pluripotent stem cell marker genes *DNMT3B, LIN28A, NANOG* and *PRDM14,* and an increase in the expression of neuronal marker genes *MAP2, RBFOX3* and *TUBB3* from day 0 to day 4 of differentiation (**Figure 2F, S7F**).

To further determine whether VLP-derived RNAs accurately captured changes in cell type-specific gene programs, we performed gene set enrichment analysis (GSEA) on the fold changes in gene expression between day 0 and day 4, using pre-defined stem cell and neuronal gene sets, as well as within KEGG-defined pathways. Indeed, we observed that genes involved in nucleic acid metabolism and stem cell division and proliferation were downregulated, while neural development-related gene expression programs were upregulated during differentiation (**Figure 2G, S7H-J**). Altogether, these results demonstrate that longitudinal applications of self-reporting can elucidate transcriptional dynamics of individual genes and gene programs within live cells undergoing differentiation.

### Epitope functionalized VLPs enable multiplexed selective self-reporting

As mammalian cells naturally exist in complex environments composed of multiple interacting cell types, we sought to selectively tailor the self-reporting readout to subpopulations of interest. While envelope proteins on the surface of virions direct viral tropism, we hypothesized that pseudotyping VLPs with engineered envelope proteins displaying unique epitope tags would enable multiplexed differential capture of VLPs produced by different cell populations (**Figure S8A**). To test this hypothesis, we engineered the vesicular stomatitis virus G (VSV-G) envelope protein with an N-terminal FLAG tag (**Figure S8B**) and co-transfected HEK293Ts with plasmids expressing this protein and Gag to pseudotype VLPs. FLAG immunoprecipitation (IP) of the cell media enabled specific, epitope-dependent isolation of VLPs (**Figure S8C**), and we observed comparable performance with HA-tagged VSV-G and HA-IP of the cell media (**Figure S8D**). RNA-seq of the IP-isolated VLPs further indicated high transcriptional signal-to-noise with this epitope-dependent isolation method versus controls (**Figure S8E**). We also observed a notable enrichment of NGS reads in intragenic and specifically exonic regions from IP-isolated VLPs (**Figure S8F**), which robustly reflected the identity of the self-reporting HEK293T cells (**Figure S8G**). Interestingly, we also found that non-viral proteins, such as the truncated nerve growth factor receptor (tNGFR), can pseudotype VLPs and result in specific purification of VLPs from the cell media when decorated with an N-terminal FLAG tag (**Figure S9A-C**). Collectively, these data suggest that VLPs pseudotyped with engineered envelope proteins enable epitope-specific isolation of VLPs from cellular media with high specificity.

To explore the potential for multiplexed selective cellular self reporting, we assessed the ability to simultaneously read out VLPs pseudotyped with distinct epitope tags corresponding to different cell types grown in co-culture. We stably introduced FLAG-tagged VSV-G into HEK293T cells and HA-tagged VSV-G into HT1080 cells as a means to uniquely identify each cell line (**Figure S10A**). We verified the epitope-specificity of our IP protocols by subjecting FLAG-tagged VLPs to HA-IP and vice versa, confirming that each protocol specifically isolated the desired epitope (**Figure S10B-C**). We then co-cultured the epitope-tagged cell lines, and following 48 hours of VLP production, subjected the cell media to FLAG- and HA-specific IPs (**Figure S10D-E**). VLP-derived RNAs isolated following FLAG and HA IPs ultimately reflected HEK293T and HT1080 cell transcriptomes, respectively (**Figure S10F**), further supporting our ability to multiplex the self-reporting readout. These data demonstrate the simultaneous and specific analysis of independent cellular self-reporting information streams from mixed populations using the multiplexed IP approach with engineered VLPs.

### Gag-RNA binding domain fusions modulate the repertoire of RNAs packaged in VLPs

We next opted to extend our VLP engineering efforts to explore whether and how the self-reported transcriptional profiles depend on the RNA binding properties of VLP components. We hypothesized that we could tailor the packaging specificities of RNAs in VLPs by altering the Gag polyprotein’s RNA binding activity with fusion to additional human RNA binding domains. To identify candidate domains, we mined human RNA binding proteins (RBPs) because they contain well-defined RNA binding domains (RBDs) known to engage diverse RNAs^49–51^. Thus, we posited that fusion of certain RBDs to the C-terminus of Gag could complement the RNA binding capacity intrinsic to the Gag NC domain (**Figure 3A**).

**Figure 3.**
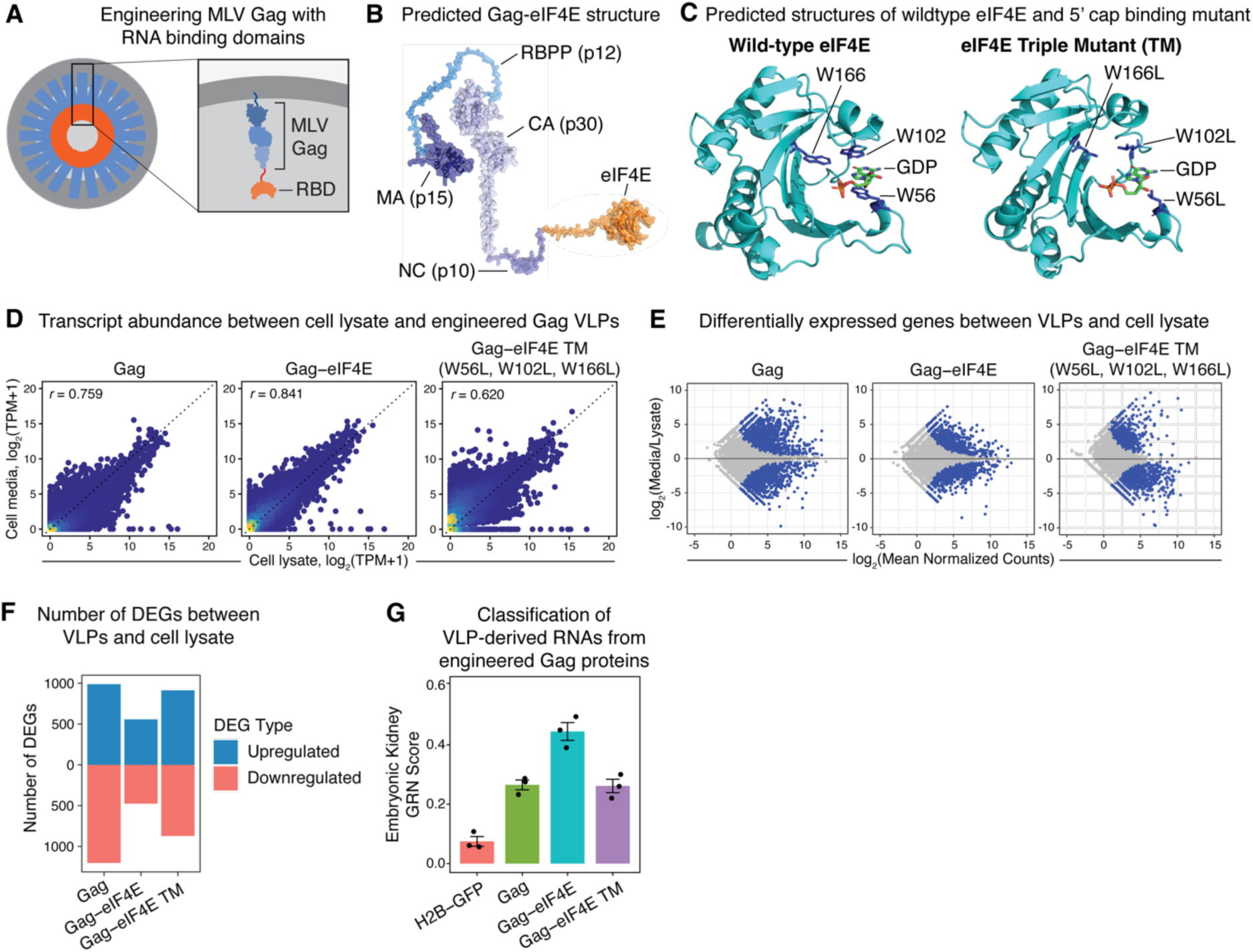
Engineering Gag with RNA binding proteins. **(A)** Schematic of VLPs derived from engineered variants of Gag fused to RNA binding proteins. **(B)** AlphaFold2-predicted structure of engineered Gag-eIF4E protein, whereby the 217-amino acid human eIF4E protein (UniProt ID: P06730) was fused to the C-terminus of MLV Gag via a 5x(GS) linker. (**C**) AlphaFold3-predicted structure of wild-type eIF4E (left) or eIF4E containing W56L, W102L, and W166L mutations (right) in complex with guanosine-5′-diphosphate (GDP). **(D)** Scatterplots of RNA transcript abundance measured from cellular lysate and VLPs purified from the cell media. HEK293T cells were transfected with plasmid vectors to express Gag, Gag-eIF4E, or Gag-eIF4E triple mutant (TM). VLPs were then purified from cell media after 48 hours. Each point represents a gene detected from RNA-seq and the values are averaged over n=3 matched replicates from cellular lysate and VLPs. Pearson correlation coefficients between cellular lysate and VLPs are presented in each plot. TPM = transcripts per million. **(E)** Bland– Altman plots (MA plots) visualizing the degree of quantitative agreement between RNA-seq measurements on cell lysate and either Gag, Gag-eIF4E, or Gag-eIF4E TM VLPs. The differences in RNA transcript abundances between cell lysate and VLPs (log2 fold change) are plotted versus the average transcript abundance measured for each gene. Statistically significant DEGs (FDR-adjusted *P* value < 0.05) are highlighted in blue. RNA count normalization was performed with DESeq2. **(F)** Quantification of the number of statistically significant up and downregulated genes detected in VLPs compared to cell lysate from (**E**). **(G)** CellNet gene regulatory network scores representing the degree to which VLP-derived RNAs reflect the GRNs in the cell lysate of the producing host cell type for various Gag VLP conditions.

We systematically assessed candidate RBDs by first curating a comprehensive list of RBPs in the human genome (from publicly available databases^52–55^) for which there exists an experimentally evaluated RNA binding motif (**Supplemental Table 3**). RNA recognition motif (RRM), hnRNP K-homology (KH), and zinc-finger (ZF) domains comprised the overwhelming majority of RBDs represented in the compendium of human RBPs (**Figure S11A**). We then utilized the experimentally-determined position weight matrices for each RBP motif to quantify the frequency and type of RNAs in the human transcriptome containing the RBP motif (**Supplemental Tables 4 and 5**). The analysis identified a core subset of RBPs containing motifs that occur frequently across non-coding and coding transcripts (**Figure S11B**). Approximately 70% of RNA transcripts contained motifs for several genes involved in poly(A)-binding, namely PABPC1 and PABPC4 (**Figure S11C**). We prioritized both RBPs for further consideration given their conserved roles across eukaryotes. Of the four N-terminal RRM domains of PABPC proteins, RRMs 1 and 2 from PABPC4 exhibit higher affinity and specificity for poly(A) RNA^56–59^. We therefore opted to fuse RRMs 1 and 2 from PABPC4 to Gag, further motivated by the high sequence conservation of these RRMs across other RRM-containing RBPs (**Figure S11D, Figure S12A**). Gag-RRM fusion proteins produced VLPs from both HEK293T and HT1080 cells (**Figure S12B**), suggesting these C-terminal modifications to Gag with RBDs are compatible with VLP assembly. Although profiling of RNAs from Gag-RRM VLPs resulted in high quality RNA-seq libraries, we did not observe a reproducible improvement in the concordance of transcript abundances from VLP-derived RNAs compared to the cell lysate (**Figure S12C-D**). This finding prompted us to consider alternative fusion proteins to enhance the packaging of RNAs into VLPs.

The majority of cellular RNAs, particularly mRNAs, contain a singular 5′ cap modification with the enzymatic addition of a *N*^7^-methylguanosine (m^7^G). eIF4E is the principal and evolutionarily conserved protein that recognizes and binds the 5′ m^7^G cap to initiate translation^60^. Thus, we fused eIF4E to the C-terminus of Gag (**Figure 3B**) and observed that VLPs could be produced from Gag-eIF4E fusion proteins (**Figure S13A**). Importantly, tryptophans 56, 102 and 166 within eIF4E are known to establish π stacking and Van der Waals interactions with m^7^G based on seminal X-ray crystal structures^61^. We modeled eIF4E bound to guanosine-5′-diphosphate (GDP) using AlphaFold3^62^, and guided by the structure, mutated the three tryptophans to aliphatic leucines to ablate the aromatic interactions with m^7^G (**Figure 3C**). We also fused this eIF4E triple mutant to Gag (termed Gag-eIF4E TM) to use as a cap binding loss-of-function control. Across independent experiments and different cell lines, we observed that RNAs derived from Gag-eIF4E VLPs exhibited higher transcriptional concordance to cell lysate than RNAs derived from wild-type Gag VLPs (**Figure 3D, Figure S13B**). We did not observe this improvement with Gag-eIF4E TM VLPs, suggesting that the higher transcriptional concordance with Gag-eIF4E could be attributed to 5′ m^7^G cap binding of RNAs (**Figure 3D**). Furthermore, we observed on average a 1.47-fold increase in the number of genes detected from Gag-eIF4E VLPs compared to wild-type Gag VLPs, and higher rates of exonic-mapped NGS reads (**Figure S13C**). This finding may point to different RNA packaging preferences of Gag-eIF4E, relative to wild-type Gag, that could enhance detection of many mature cytosolic transcripts. To further explore RNA packaging biases, we utilized previously published RiboSeq data in HEK293T cells^63^ to identify genes undergoing active translation. We observed a global enrichment of RiboSeq-detected genes within Gag-eIF4E VLPs compared to wild-type Gag VLPs (**Figure S13D**). These data are consistent with the conserved role of eIF4E in translation initiation on spliced, cytoplasmic RNAs and indicate that Gag-RBP fusions can alter the global repertoire of RNAs packaged within VLPs. Despite these global differences in exported transcriptional profiles, expression of Gag-eIF4E, similar to that of wild-type Gag, was minimally perturbative transcriptionally to cells (**Figure S3F, S13E**).

We next performed differential gene expression analysis to quantify transcriptional differences between VLP-derived RNAs and cell lysate measurements for all our engineered Gag variants (**Figure 3E**). Strikingly, we quantified approximately half as many DEGs between VLPs and cell lysate with Gag-eIF4E (557 genes significantly upregulated, and 476 significantly downregulated), compared to wild-type Gag (988 genes significantly upregulated, and 1,203 significantly downregulated) (**Figure 3F**), suggesting that RNAs exported in Gag-eIF4E VLPs better matched the profile represented in cell lysate measurements. Again, this improvement was largely abrogated with Gag-eIF4E TM (913 genes significantly upregulated, and 872 significantly downregulated), providing evidence that the difference is specifically attributable to the cap-binding activity of eIF4E. The reduction in DEGs appeared most concentrated in low-to-mid expressed genes (**Figure 3E**). The higher transcriptional similarity to cell lysate was also associated with improved CellNet classification of self-reported RNAs to cellular identity of the host transcriptome from Gag-eIF4E VLPs compared to wild-type Gag VLPs (*P* = 0.0122, **Figure 3G, Figure S13F**). Taken together, our data suggest that Gag protein engineering can alter and improve the representation of RNA molecules in VLPs for transcriptional profiling applications.

### VLPs support export of RNA barcode transcript libraries

The observation that RBPs can modulate the repertoire of RNAs packaged in VLPs inspired us to further engineer the self-reporting system to achieve targeted export of transcript populations of interest, such as nucleic acid barcode libraries for lineage tracing or CRISPR screening applications. Specifically, we sought to perform directed export of CROPseq synthetic transcripts to enable self-reporting readout of barcode libraries^64^. To direct the specificity of RNA export toward desired transcripts, we selected a minimal hairpin aptamer, which selectively binds the MS2 bacteriophage coat protein (MCP)^36^. We introduced 24 tandem repeats of MS2 hairpins into the CROPseq vector, since it can serve as a generalized expression system to encode nucleic acid barcodes within a polyadenylated RNA transcript (**Figure S14A**). We equivalently fused MCP to the C-terminus of Gag, which we validated could produce VLPs (**Figure S14B-C**). Lentivirus-mediated introduction of the CROPseq vector and Gag-MCP into HEK293T cells resulted in a 12-fold enrichment of CROPseq barcode transcripts within VLPs compared to cell lysate (**Figure S14D-E**). CROPseq barcode transcripts were also 10-fold more abundant in Gag-MCP VLPs compared to wild-type Gag VLPs (**Figure S14F-G**), confirming the utility of hairpin aptamers for preferentially packaging synthetic RNA transcripts as previously reported^36^. We then leveraged this system and designed molecular dialout strategies to read out self-reported barcode libraries from a population of cells (**Figure S14H**). We constructed a pooled library of barcodes (**Supplemental Table 6**), which we cloned into the CROPseq vector and introduced into HEK293T cells via low MOI lentivirus infection without significant biases in the representation of individual barcodes (**Figure S14I-J**). Following introduction of either wild-type Gag or Gag-MCP into the barcoded population and subsequent production of VLPs, we dialed out CROPseq barcode transcripts from VLPs and quantified the abundance by NGS in the manner of a pooled genetic screen. Gag-MCP VLPs enabled barcode abundance quantification that better matched ground truth abundance measured from genomic DNA than wild-type Gag VLPs (Pearson *r* = 0.822 versus 0.477, **Figure S14K**). These data support that the self-reporting system can be tailored to read out specific RNA transcripts and barcode libraries with high accuracy.

## DISCUSSION

In this work, we developed ‘self-reporting,’ a synthetic export system in which cellular RNAs are packaged into VLPs for non-destructive, longitudinal transcriptome-wide profiling of dynamic biological processes from living cells. Compared to traditional RNA-seq, live-cell RNA-seq offers a number of key advantages. First, self-reporting allows a living cell sample to be monitored over time without lysing the cells themselves and enables downstream studies on the same living biological sample that remains after collection of VLPs for RNA-seq. This non-destructive sampling is especially useful for non-dividing cells, including neurons, myocytes, and adipocytes. Additionally, biological models, such as 3D organoid cultures, and processes, such as cellular differentiation, that exhibit strong instance-to-instance variability are particularly poised to benefit from analysis of dynamics over serial measurements from a single instance. In addition to transcriptome-wide profiling, we showed that self-reporting can also enable detection of specific transcript populations of interest, such as nucleic acid barcodes and sgRNAs for CRISPR screening. We envision this platform to be useful for monitoring population dynamics, including in live-cell pooled genetic perturbation screens.

However, there are important considerations of our current self-reporting protocol. The signal from VLP-derived RNAs collected at any time point is an integration over the sampling duration, rather than a snapshot at a single time point. Additionally, VLP sampling requires a media change at each time point, which may adversely affect cell fitness. As a result, self-reporting has an inherent tradeoff between time resolution or sensitivity and alteration of the environmental conditions experienced by the cells. We reason that this issue could be mitigated by replacing sampled media with conditioned culture medium. Furthermore, RNAs exported in VLPs represent a fraction of the RNAs that are present in the cells, which may result in lower statistical power for differential gene expression calls from self-reported relative to lysate measurements. Consistent with this reasoning, we observed that gene expression changes detected in self-reported RNA were strongly but not perfectly correlated to those in cell lysates. In this study, we deliberately expressed the Gag protein at low levels to minimize perturbation of the cells. However, the expression level of Gag could be increased constitutively or transiently to increase VLP export and improve sensitivity for key dynamic transitions occurring in a specific time window. We also demonstrated the ability to improve the fidelity of the transcriptional profiles recovered from VLPs by fusing Gag to RNA binding proteins. Further engineering of Gag to enhance RNA recognition and packaging into VLPs is likely to increase the sensitivity of the self-reporting readout.

We envision that future applications of VLP-based live-cell self-reporting could ultimately encompass targeted longitudinal transcriptomic measurements of single cells, tissues, and organs within their natural physiological and spatial context. For example, the expression of self-reporting machinery could be driven by cell-type-specific promoters in transgenic animals to monitor the transcriptomes of living target cells *in situ*. In addition, although we demonstrated that we can detect self-reported RNAs from single cells, future improvements to RNA export efficiency may enhance sensitivity and time resolution, thereby supporting more robust single-live-cell transcriptomics experiments. We also envision that self-reporting may be adapted for non-destructive proteomic and metabolomic measurements from VLP cargo other than RNA. Altogether, our work establishes cellular self-reporting with engineered VLPs as an attractive tool for live-cell transcriptomics, enabling the study of dynamic biological processes across diverse cell types.

## ACKNOWLEDGMENTS

We thank A. Wassie, D. Ter-Ovanesyan, A. Dixit, O. Parnas, D. Feldman, T. Kulesa, Z. Bloom-Ackermann, C. Ackerman, F. Zhang, F. Chen, E. Boyden, D. Lauffenburger, G. Church, J. Collins, and members of the Daley and Blainey labs for helpful discussions and technical advice. We thank E. Kowal and W. Salmon for assistance with electron microscopy. We thank the Church lab at Harvard Medical School for contributing the PGP1 iNGN cell line. We thank F. McCarthy, E. Botelho, and R. Dertinger for support with grant administration and laboratory operations. We thank F. Chen, X. Wang, A. Hansen, A. Shalek, B. Deverman, K. Caetano-Anolles, R. Majovski, and P. Chen for feedback during the preparation of this manuscript. We thank the NIH Single Cell Analysis Program (SCAP) and InnoCentive, Inc. for creating the *Follow that Cell* Challenge and the other program participants for creating a scientific community around longitudinal biological analyses.

## AUTHOR CONTRIBUTIONS

J.B., M.N., and P.C.B. contributed to conceptualization of the project. J.B. performed initial proof-of-concept experiments. M.N., J.B., A.L., F.T., and J.Y.H. designed and performed additional experiments. M.N., J.B., and A.L. analyzed data. M.N., A.L., and P.C.B. prepared the manuscript with input from all authors. L.G.G., G.Q.D., and P.C.B. supervised the research. G.Q.D. and P.C.B. provided funding.

## SUPPLEMENTAL FIGURES

**Figure S1.**
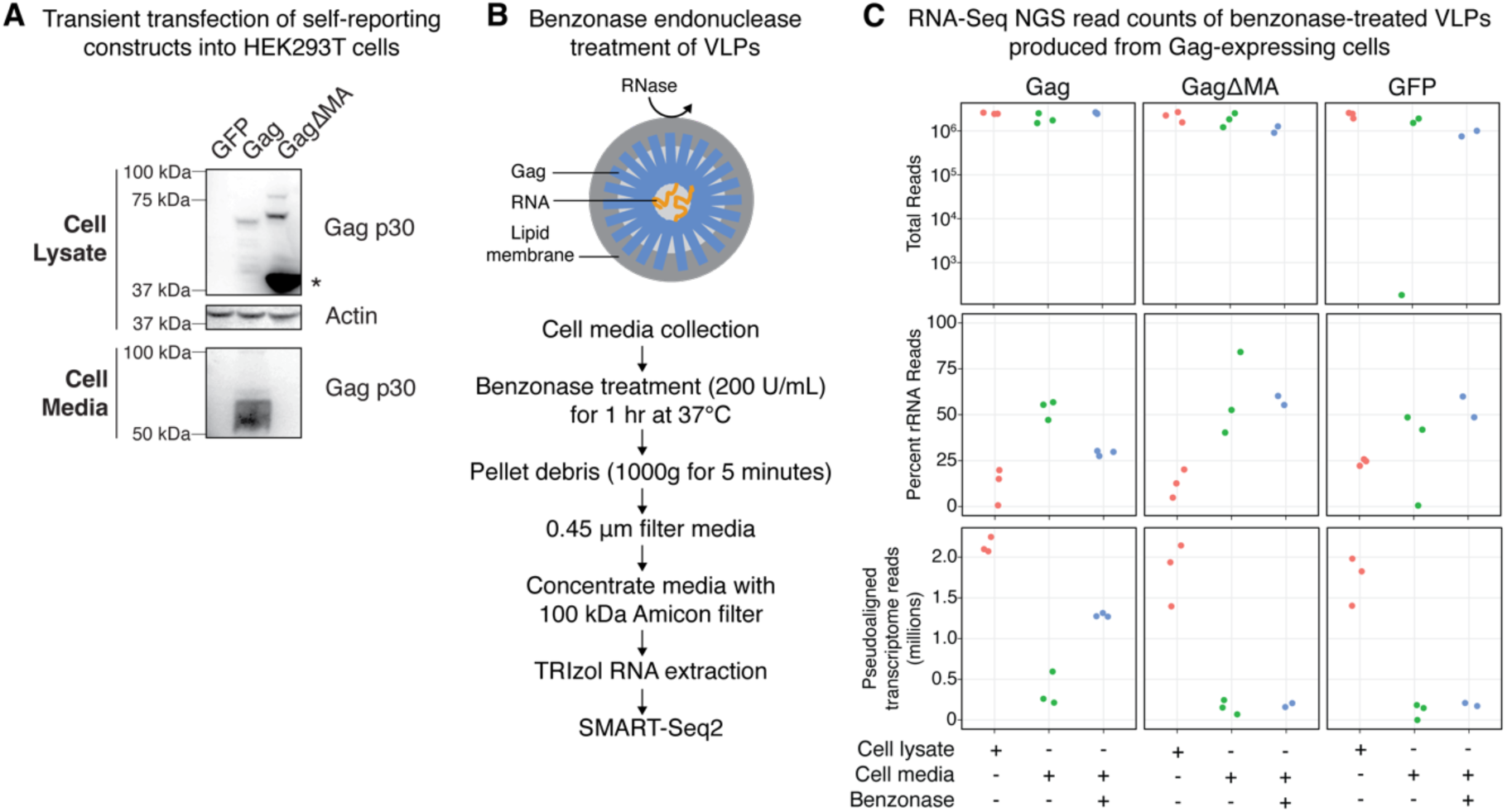
Nuclease protection of cellular RNAs packaged in VLPs. **(A)** Western-blot verification of VLP production from HEK293T cells transfected with either GFP, Gag, or GagΔMA expression vectors. Protein from cells and culture media was extracted 48 hours post-transfection and immunoblotted for MLV Gag p30 in both cell lysate and media fractions, as well as for actin in cell lysate as a loading control. Expected molecular weights: 62.07 kDa for Gag and 38.74 kDa for GagΔMA (monomer is marked with (*) and heavier bands are concatemers). **(B)** Experimental protocol to test the hypothesis that RNAs packaged in VLPs are protected from nuclease degradation. Cell media from Gag, GagΔMA, and GFP-only expressing cells were treated with benzonase endonuclease, a non-specific RNA/DNA nuclease. VLPs were purified from the cell media, after which RNA-seq libraries were prepared via SMART-Seq2 and sequenced with next generation sequencing (NGS) to quantify transcriptome-wide gene expression. **(C)** Quantification of RNA-seq NGS reads on cell media +/− benzonase treatment from Gag, GagΔMA, and GFP-only expressing cells, highlighting the total sequencing depth of each RNA-seq library (top), the proportion of rRNA reads in each library (middle), and the number of human transcriptome aligned reads per library (bottom). Cell lysate samples are included as SMART-Seq2 positive controls. Each condition is represented by n=3 replicates.

**Figure S2.**
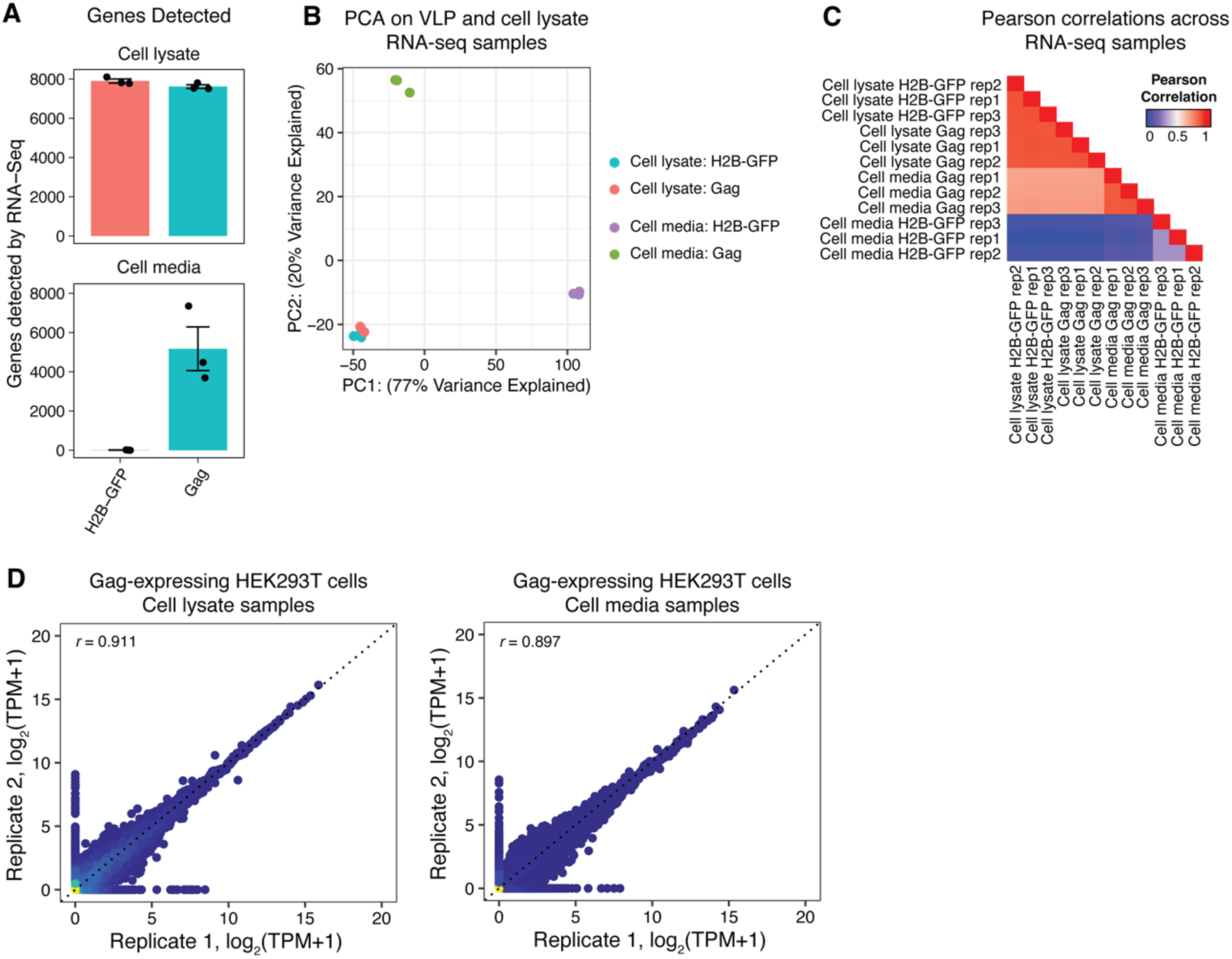
Characterization of cells transiently transfected with MLV Gag. **(A)** Quantification of the number of genes detected by RNA-seq from cell lysate and cell media samples from HEK293T cells transfected with H2B-GFP or Gag expression vectors. Each point represents an independent transfection replicate. (**B**) Two-dimensional PCA scores plot of cell lysate and cell media transcriptional profiles from H2B-GFP or Gag-expressing HEK293T cell. **(C)** Pearson correlations of measured transcriptional abundances across all RNA-seq samples. **(D)** Concordance of transcript abundances between two transfection replicates of cell lysate (left) and cell media (right). Pearson correlation coefficients are noted within each plot.

**Figure S3.**
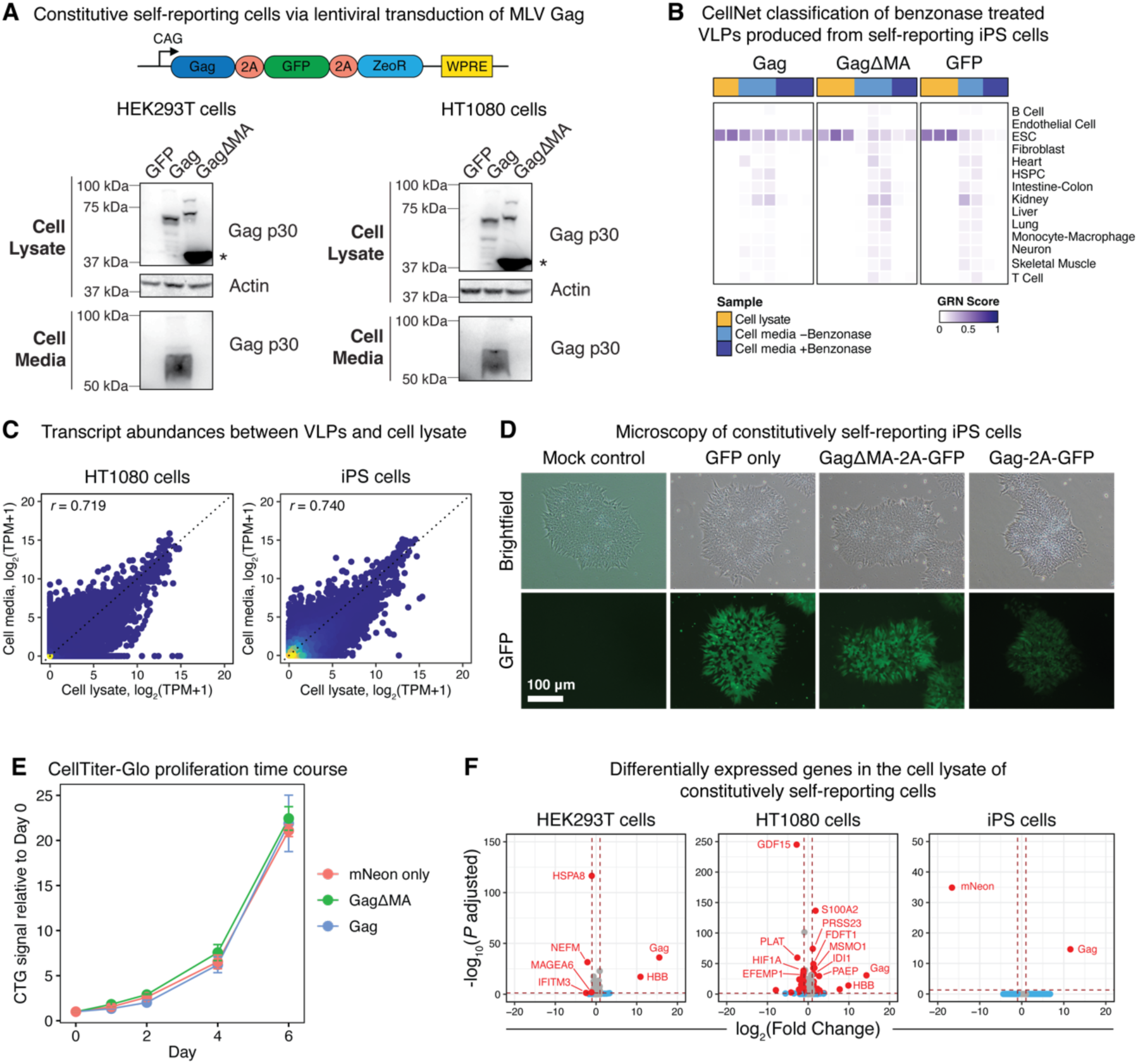
Characterization of cells engineered to constitutively express MLV Gag. **(A)** Western-blot verification of VLP production from HEK293T and HT1080 cells lentivirally transduced with either GFP, Gag, or GagΔMA constitutive expression vectors (schematically depicted). Cells were incubated for 48 hours to produce VLPs and then protein was extracted from cells and the culture media. Samples were assayed for MLV Gag p30 in both cell lysate and media fractions, as well as for actin in cell lysate as a loading control. Expected molecular weights: 62.07 kDa for Gag and 38.74 kDa for GagΔMA (monomer is marked with (*) and heavier bands are concatemers). **(B)** CellNet GRN classification analysis of RNA-seq libraries of cell lysate and cell media +/− benzonase treatment produced from iPS cells constitutively expressing GFP, GagΔMA, or Gag. iPS cell media was collected and processed following 48 hours for VLP production. **(C)** Scatterplot of RNA transcript abundances measured from cellular lysate and Gag VLPs for HT1080 cell (left) and iPS cells (right) stably expressing Gag. Each point represents a gene detected from RNA-seq and the values are averaged over n=3 biological replicates. Pearson correlation coefficients between cellular lysate and VLPs are noted in each plot. TPM = transcripts per million. **(D)** Brightfield and fluorescence microscopy of iPS constitutively expressing GFP only, GagΔMA, or Gag. **(E)** Cell proliferation time-course of cells constitutively expressing mNeon alone, GagΔMA, or Gag measured by a CellTiter-Glo assay. **(F)** Volcano plots of differentially expressed genes in the cell lysate between control and Gag-expressing HEK293T, HT1080, and iPS cells. Positive log2(fold change) values are reflective of upregulated genes in Gag-expressing cells. Significant differentially expressed genes are defined as log2(fold change) > 1 or log2(fold change) < −1 and an FDR-adjusted *P* value below 0.05.

**Figure S4.**
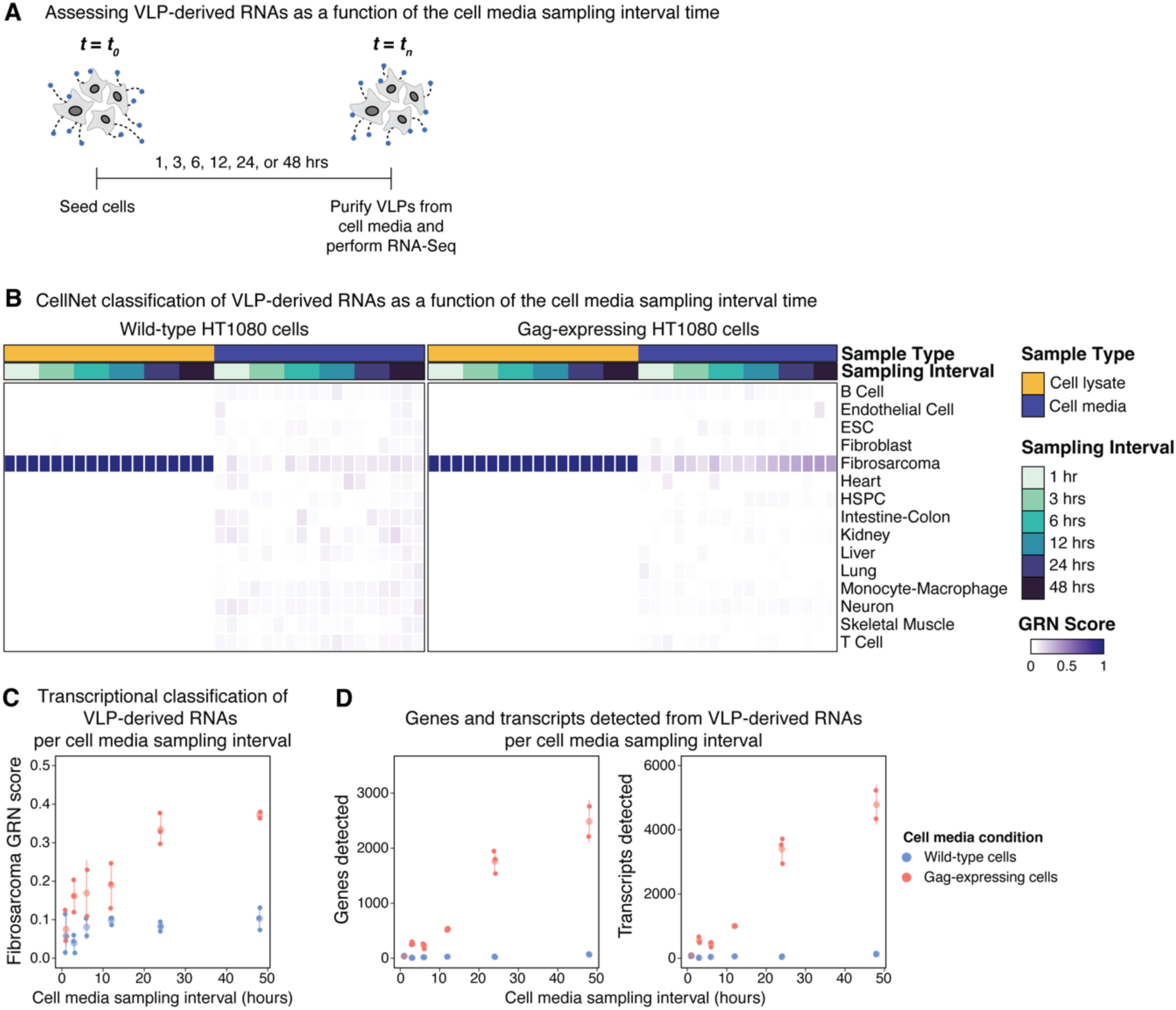
Temporal kinetics of RNA export in VLPs. **(A)** Experimental schematic to characterize self-reported RNAs in VLPs as a function of the amount of time allowed for VLP production (hereto referred to as the cell media sampling interval). **(B)** CellNet GRN classification analysis on cell lysate and cell media from constitutively Gag-expressing and wild-type HT1080 cells. The cells were incubated for 1, 3, 6, 12, 24, or 48 hours (n=3 replicates per time-point and cell line), after which RNA-seq libraries were prepared on VLP-purified cell media and the associated cell lysate. The CellNet classifier was retrained with the inclusion of HT1080 cell lysate samples to incorporate a fibrosarcoma gene regulatory network. **(C)** Fibrosarcoma GRN scores on the cell media of wild-type and Gag-expressing HT1080 cells as a function of the cell media sampling interval time. **(D)** The number of genes (left) and transcripts (right) detected via Kallisto pseudoalignment to the hg19 transcriptome from the cell media of wild-type and Gag-expressing HT1080 cells as a function of the cell media sampling interval time.

**Figure S5.**
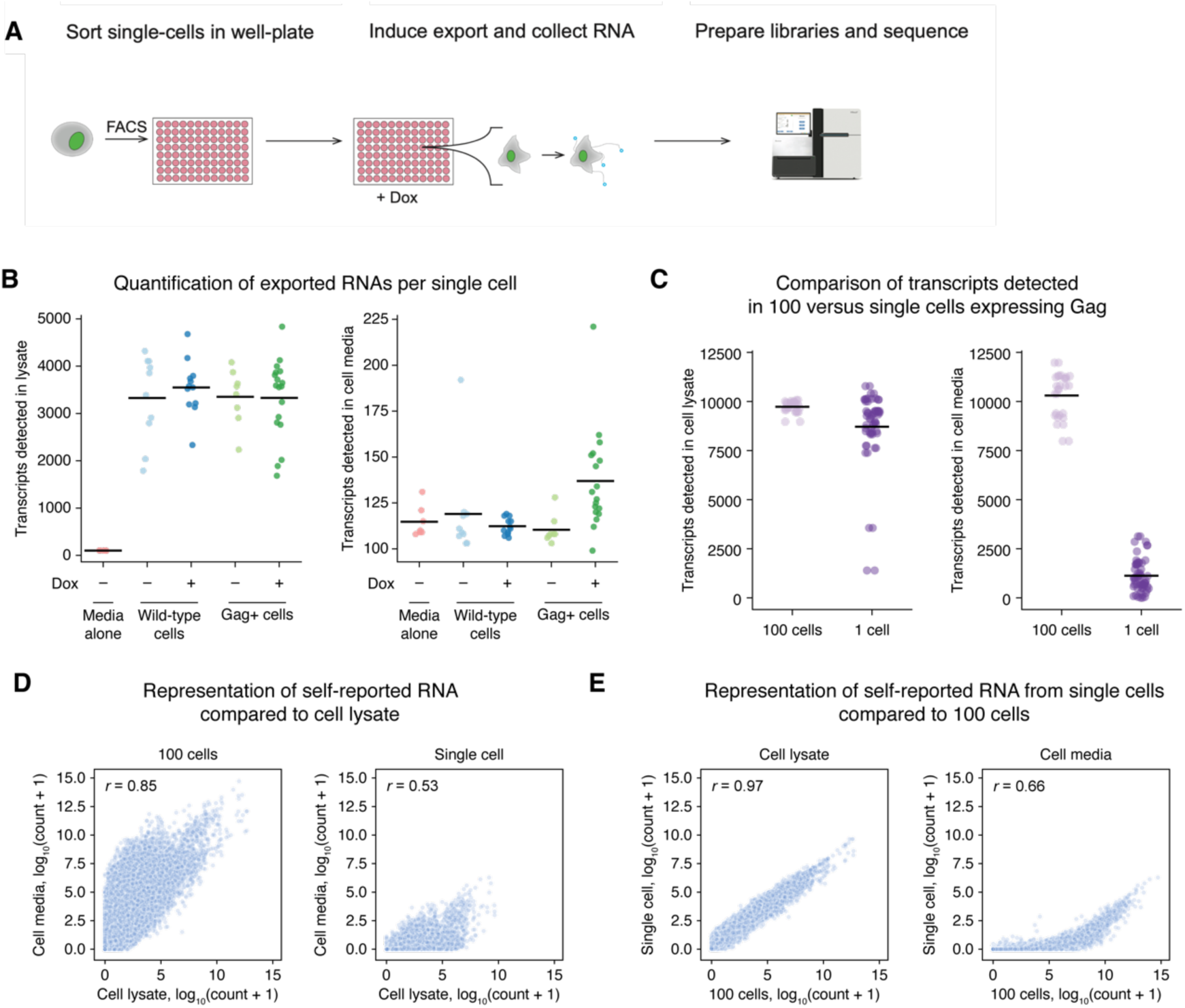
Characterization of RNA export rates from self-reporting single cells. **(A)** Experimental schematic to quantify RNA export rates from single cells over 24 hours of VLP production. Single HEK293T cells engineered with dox-inducible Gag were FACS sorted into 96-well plates and cultured for 24 hours with 1 µg/mL dox. Brightfield microscopy was performed immediately prior to and following cell media collection to identify wells containing a viable cell. RNA-seq libraries were then prepared from cell media and cell lysate via SMART-Seq2 and sequenced to quantify transcriptome-wide gene expression. Samples were retained for downstream analysis if cell lysates had greater than 1,000 transcripts detected. **(B)** Number of transcripts detected in the cell lysate (left) and cell media (right) of single cells. **(C)** Comparison of the number of transcripts detected in the cell lysate (left) and cell media (right) of 100 or single HT1080 cells stably and constitutively expressing Gag. Cells were sorted into 384-well plates and cultured for 24 hours before cell media collection and cell lysis for RNA-seq. **(D)** Transcript abundance concordance between cell lysates and cell media of 100 cells (left) and single cells (right) constitutively expressing Gag. Transcript counts were obtained from alignment using STAR. Pearson correlation coefficients *r* are presented in each plot. **(E)** Transcript abundance concordance between 100 cells and single cells measured from cell lysate (left) and cell media (right). Pearson correlation coefficients *r* are presented in each plot.

**Figure S6.**
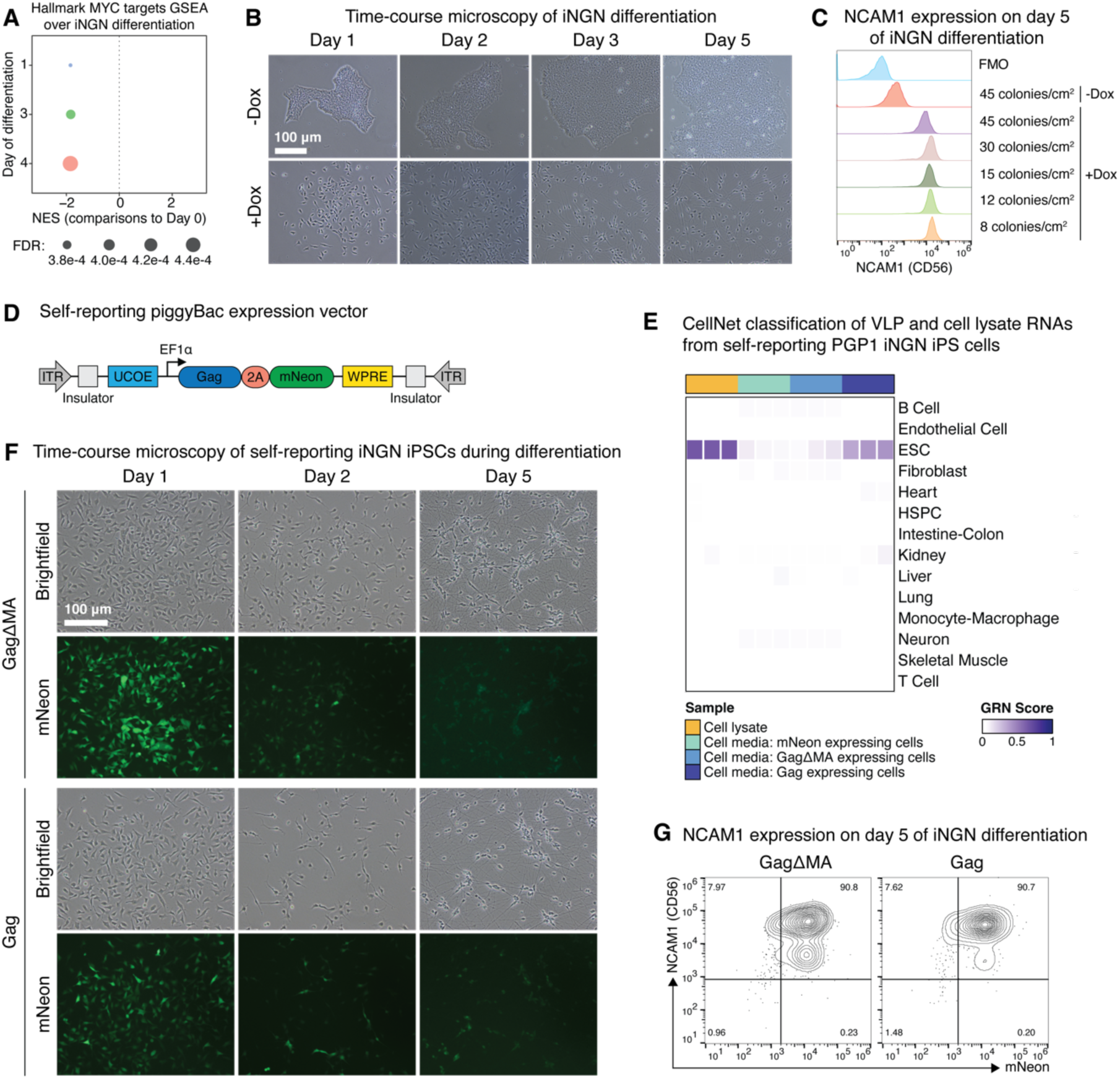
Differentiation from self-reporting PGP1 iNGN iPS cells to bipolar-like neurons. **(A)** GSEA for the Hallmark MYC targets gene set on temporally differentially expressed genes over the course of iNGN differentiation (RNA-seq data reanalyzed from Busskamp, *et al.* 2014, PMID: 25403753). Differential expression at each time point was determined relative to day 0 of the differentiation. Genes were ranked based on descending log2(fold change) from DESeq2 and negative NES scores reflect gene set depletion in a sample relative to day 0. *P* values were FDR corrected for multiple hypothesis testing. **(B)** Brightfield microscopy of neural differentiation over 5 days from PGP1 iNGN iPS cells cultured with 1 µg/mL dox (to induce expression of Neurogenin-1 and Neurogenin-2 transcription factors) or in the absence of dox (negative control). **(C)** Flow cytometry quantification of NCAM1 cell surface expression following 5 days of differentiation from PGP1 iNGN cells plated at various densities on day 0. **(D)** piggyBac expression vector to constitutively express MLV Gag in PGP1 iNGN iPS cells. **(E)** CellNet classification of cellular lysate and VLP-derived RNAs from PGP1 iNGN iPS cells engineered to constitutively express MLV Gag, GagΔMA, or mNeon only. The GRN score represents the degree to which the sample recapitulates the GRN of known cell types within the CellNet classifier. n=3 biological replicates per condition. **(F)** Microscopy of neural differentiation over 5 days from Gag or GagΔMA expressing PGP1 iNGN iPS cells cultured with 1 µg/mL dox. **(G)** Flow cytometry quantification of NCAM1 and mNeon expression following 5 days of differentiation from Gag or GagΔMA expressing PGP1 iNGN iPS cells.

**Figure S7.**
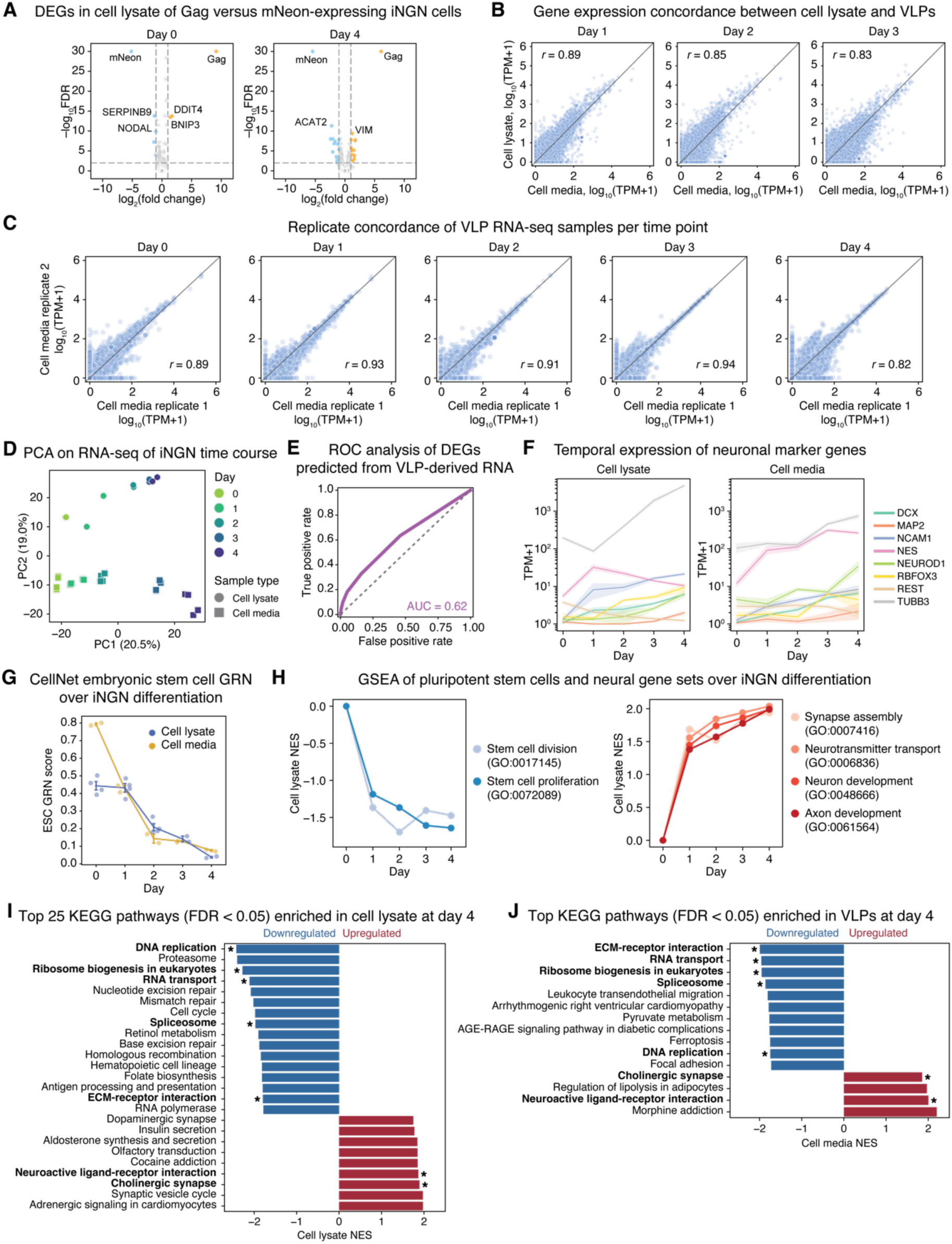
Self-reporting of transcriptional dynamics during neuronal differentiation of iNGN cells. **(A)** Volcano plots of differentially expressed genes (DEGs) between Gag-expressing and mNeon-expressing PGP1 iNGN cells at days 0 (left) and 4 (right) of neuronal differentiation. DEGs were defined as genes with log2(fold change) > 1 or log2(fold change) < −1 and false discovery rate (FDR)-adjusted *P* value below 0.01. **(B)** Gene expression concordance between cellular lysates and VLPs at days 1, 2, and 3 of iNGN differentiation. Log-transformed transcripts per million (TPM) were averaged across replicates (n=2 for cellular lysates, n=4 for VLPs). Pearson correlation coefficients *r* are presented in each plot. **(C)** Gene expression concordance between 2 biological replicates of RNA obtained from VLPs from day 0 to day 4. Pearson correlation coefficients *r* are presented in each plot. **(D)** Two-dimensional PCA scores plot of transcriptional profiles obtained from cellular lysates (circle) and VLPs (square) between days 0 and 4. **(E)** ROC curve for predicting differentially expressed genes in cell lysate based on log2 fold changes in gene expression detected in VLPs between day 0 and day 4 of iNGN differentiation (area under the curve, AUC = 0.62). **(F)** Gene expression levels for neural markers over 4 days of differentiation in cellular lysate (top, mean ± SEM of n=2 replicates) and VLPs (bottom, mean ± SEM of n=4 replicates). **(G)** Line plot of ESC gene regulatory network (GRN) scores obtained from CellNet. CellNet was performed on RNA derived from cellular lysate and VLPs at each time point to assess the degree to which samples classified as embryonic stem cells (ESCs). **(H)** Line plots of normalized enrichment scores (NES) for select gene ontology (GO) terms describing stem cell states (left) and neuron development (right) across 4 days of differentiation. GSEA was performed on ranked log2 fold changes in gene expression between each time point to day 0 from cellular lysate samples. **(I-J)** Bar plots quantifying GSEA NESs for KEGG pathways with FDR-adjusted *P* values < 0.05 in cellular lysate **(I**, top 25 pathways) and VLPs **(J)**. GSEA was performed on ranked log2 fold changes in gene expression between days 0 and 4. Asterisks indicate terms that are shared between cell lysate and VLP measurements.

**Figure S8.**
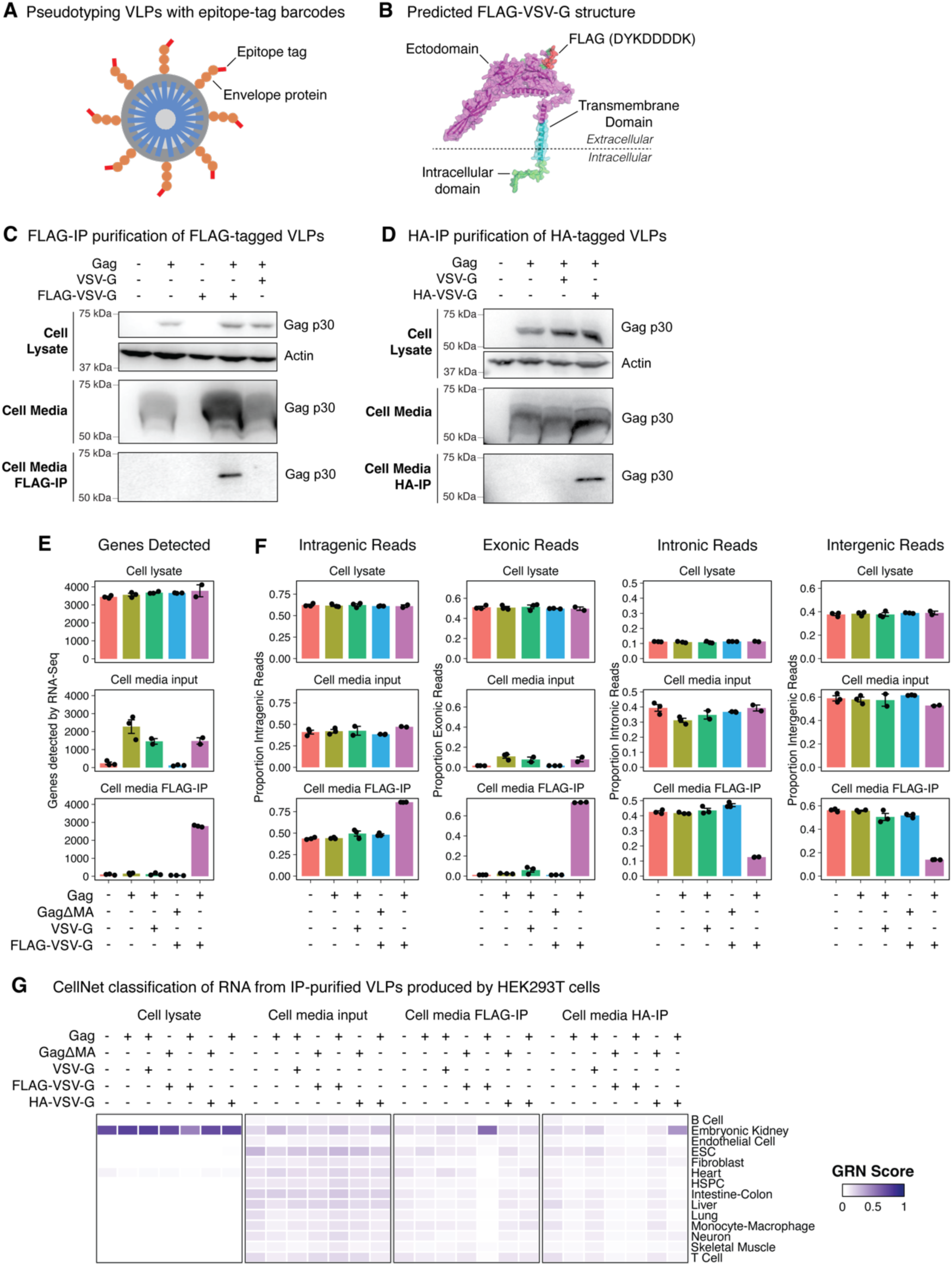
Pseudotyping VLPs with epitope-tag barcodes. **(A)** Experimental schematic to pseudotype VLPs with engineered envelope proteins containing epitope-tag barcodes. **(B)** AlphaFold2-predicted structure of VSV-G containing a FLAG tag inserted within the ectodomain following the signal peptide. **(C-D)** Western-blot validation of immunoprecipitation-based isolation from cell media of VLPs pseudotyped with epitope-tagged VSV-G proteins. HEK293T cells were transfected with plasmids to express Gag, wild-type VSV-G, and either FLAG-tagged VSV-G **(C)** or HA-tagged VSV-G **(D)**. The cell lysate and media were collected 48 hours post-transfection and the cell media was subject to FLAG-IP **(C)** or HA-IP **(D)**. Protein was extracted from all samples and assayed for MLV Gag p30 in both cell lysate and media fractions, as well as for actin in cell lysate as a loading control. The expected molecular weight of MLV Gag is 62.07 kDa. **(E)** Quantification of the number of genes detected by RNA-seq in the cell lysate, cell media, and FLAG-IP on cell media from HEK293T cells transfected with plasmids to express GagΔMA, Gag, wild-type VSV-G, or FLAG-tagged VSV-G. **(F)** Proportion of mapped RNA-seq reads to intragenic, exonic, intronic, or intergenic regions for the same samples represented in **(E)**. **(G)** CellNet GRN classification analysis on RNA-seq libraries of IP-purified VLPs pseudotyped with FLAG or HA-tagged VSV-G. The CellNet classifier was retrained with the inclusion of HEK293T cell lysate samples to incorporate an embryonic kidney gene regulatory network.

**Figure S9.**
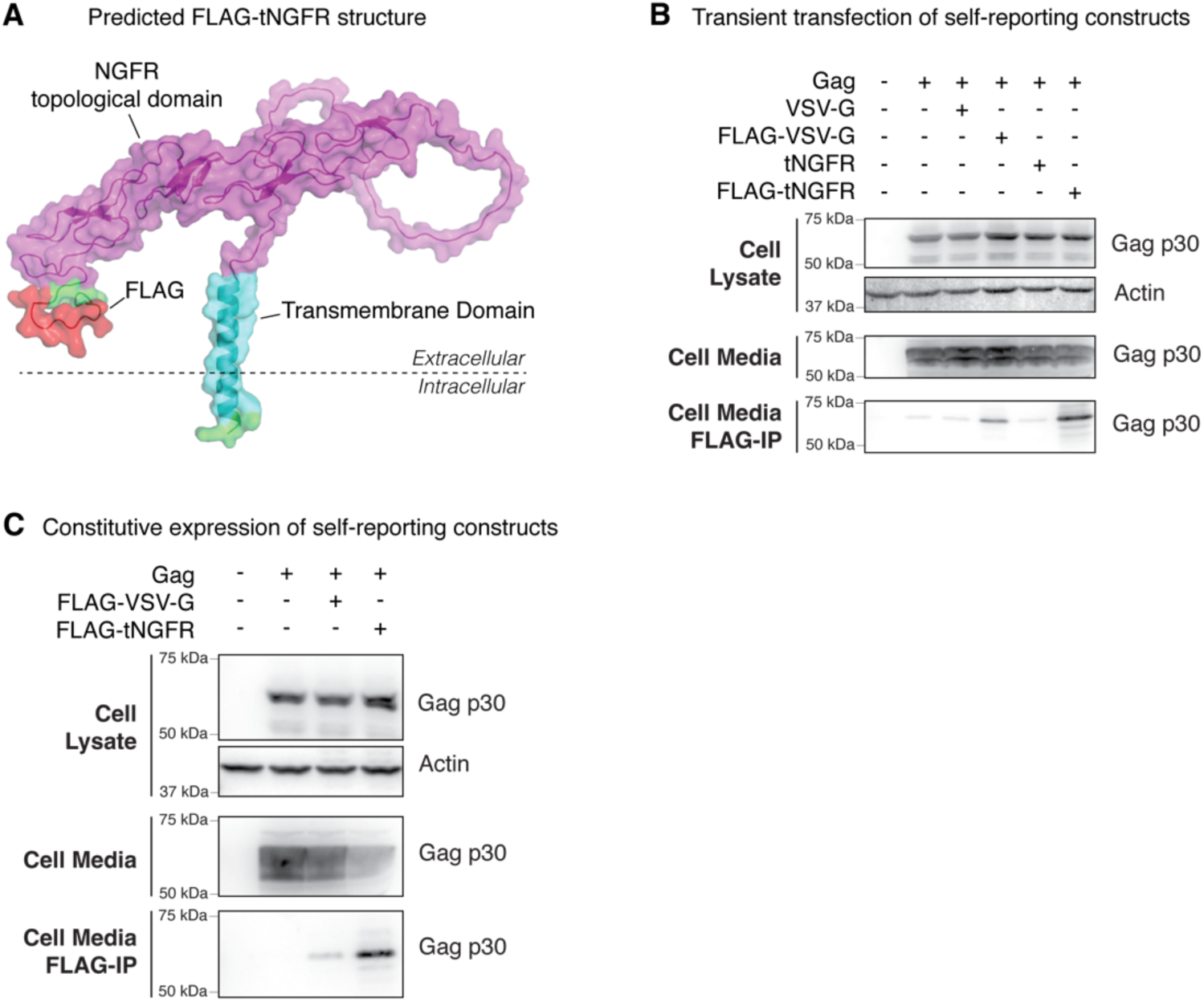
Non-viral surface proteins can pseudotype VLPs. **(A)** AlphaFold2-predicted structure of the truncated nerve growth factor receptor (tNGFR) containing a FLAG tag inserted within the NGFR topological domain following the signal peptide. **(B)** Western-blot validation of immunoprecipitation-based isolation of FLAG-tagged tNGFR pseudotyped VLPs from cellular media. HEK293T cells were transfected with plasmids to express Gag, wild-type VSV-G, FLAG-tagged VSV-G, tNGFR, or FLAG-tagged tNGFR. The cell lysate and media were collected 48 hours post-transfection and the cell media was subject to FLAG-IP. Protein was extracted from all samples and assayed for MLV Gag p30 in both cell lysate and media fractions, as well as for actin in cell lysate as a loading control. The expected molecular weight of MLV Gag is 62.07 kDa. **(C)** Equivalent experiment as **(B)** but HEK293T cells were engineered via piggyBac nucleofection to constitutively express FLAG-tagged VSV-G or FLAG-tagged tNGFR.

**Figure S10.**
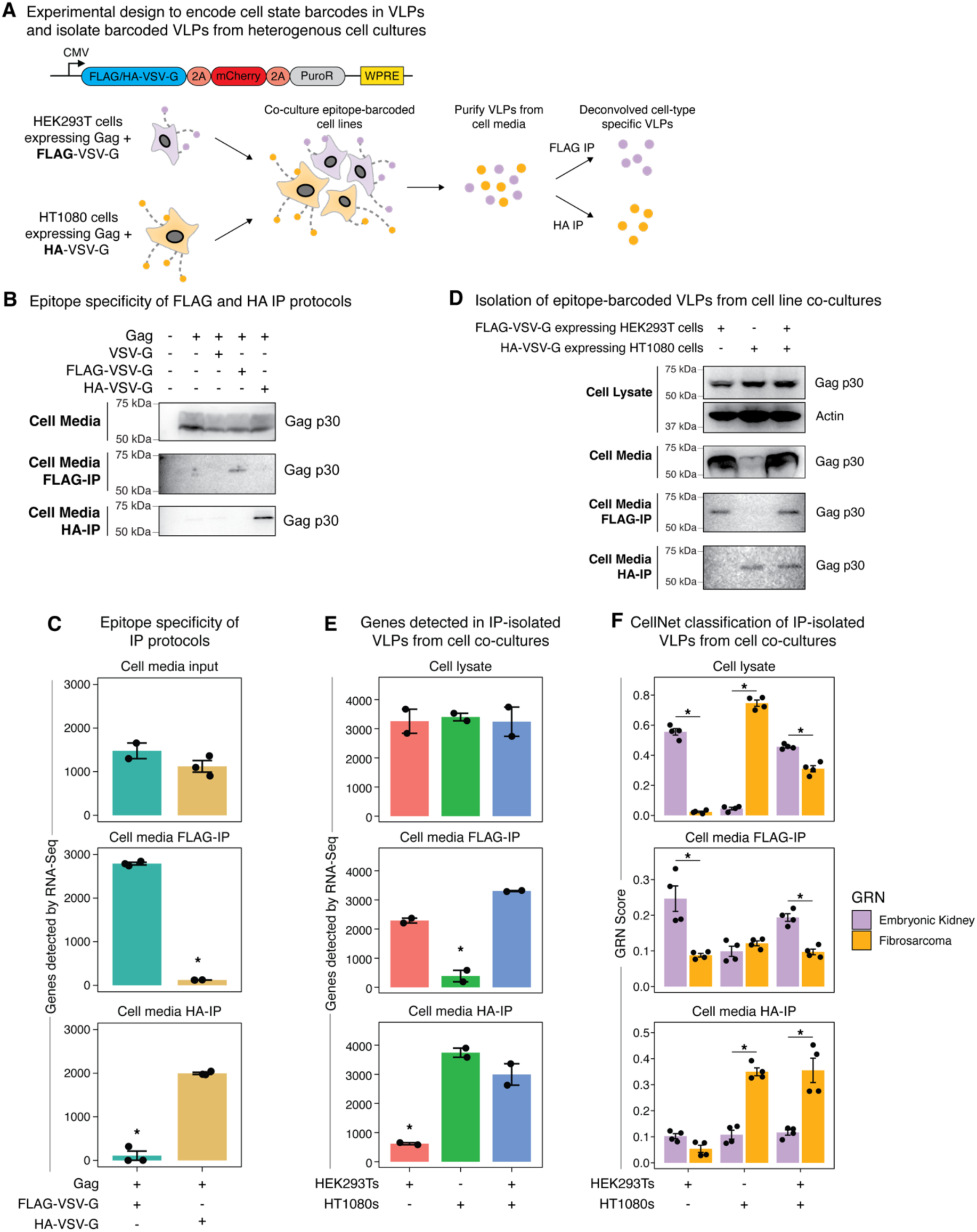
Deconvolution of epitope-tag barcoded VLPs from cell line co-cultures. **(A)** Experimental design to isolate cell-type specific VLPs from cell line co-cultures via immunoprecipitation of epitope-tag barcodes. Gag-expressing HEK293T and HT1080 cells were lentivirally transduced with FLAG or HA-tagged VSV-G expression vectors (schematically depicted), respectively. Self-reporting HEK293T and HT1080 cells were pooled and cultured together for 48 hours to allow for VLP export into the cell media. The cell media from the co-cultures was then subjected to FLAG and HA IPs to isolate HEK293T and HT1080 VLPs, respectively. **(B)** Western-blot to verify the epitope-specificity of FLAG and HA IP protocols. FLAG and HA-tagged VSV-G pseudotyped VLPs were produced via transient transfection into HEK293T cells and then processed via FLAG and HA IPs. All samples were assayed for MLV Gag p30. The expected molecular weight of MLV Gag is 62.07 kDa. **(C)** Quantification of the number of genes detected by RNA-seq from FLAG and HA-tagged VSV-G pseudotyped VLPs subjected to both FLAG and HA IPs. **(D)** Western-blot validation of cell type-specific isolation of VLPs from cell line co-cultures via epitope-tag based IPs on the cell media. **(E)** Quantification of the number of genes detected by RNA-seq from the samples in **(D)**. **(F)** CellNet classification scores for embryonic kidney and fibrosarcoma GRNs from the samples in **(D)**. Asterisks indicate an adjusted *P* value < 0.05, computed with a two-tailed Wilcoxon rank sum test and corrected multiple hypothesis testing.

**Figure S11.**
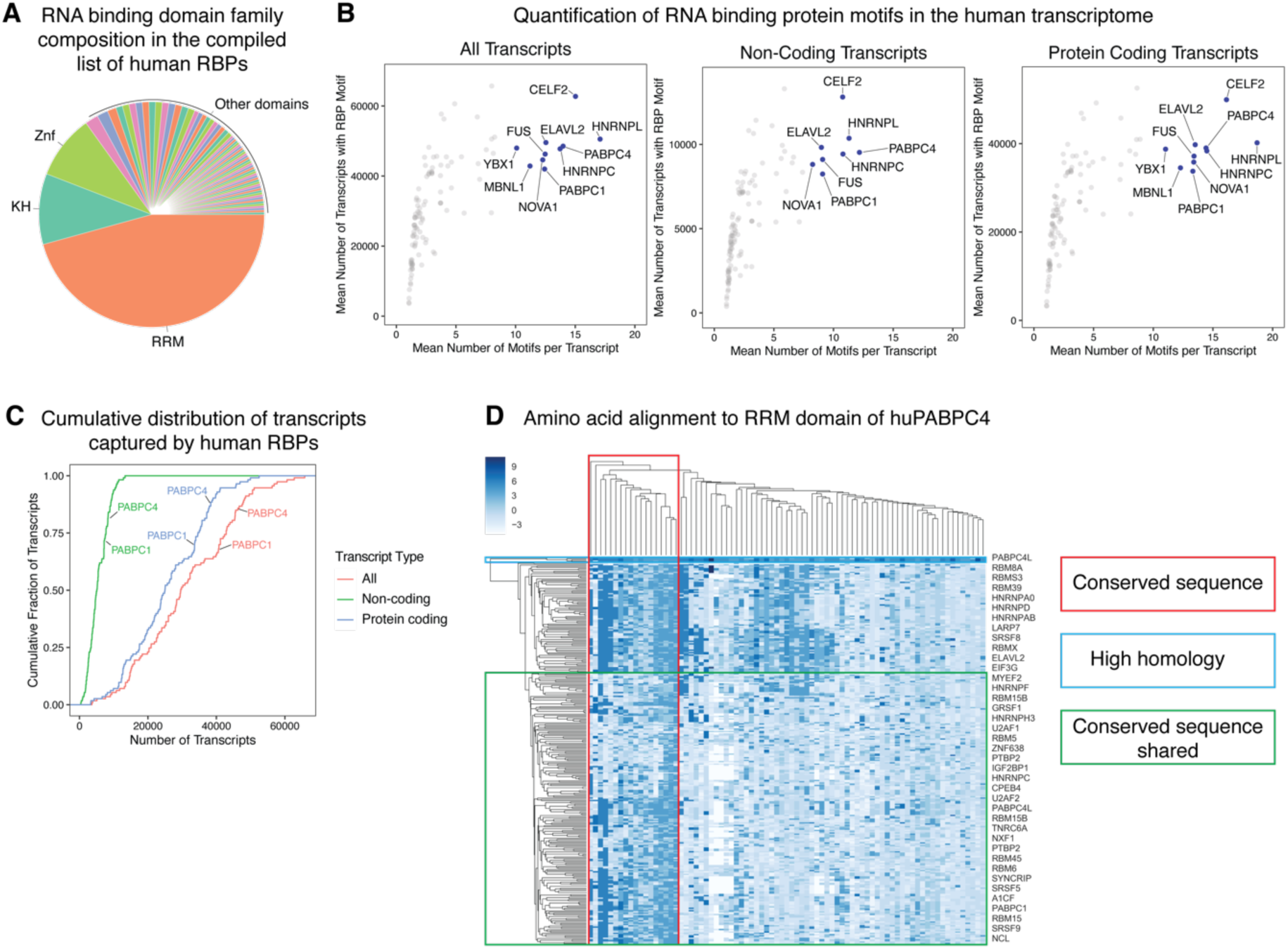
Quantification and characterization of RNA binding protein motifs in the human transcriptome. **(A)** Proportion of RNA binding domain families represented within the comprehensive curated set of human RBPs. **(B)** Quantification of the number of RNA transcripts containing RBP motifs versus the number of motifs per transcript for all transcripts in the human transcriptome (left, 70,863 transcripts), non-coding transcripts (middle, 16,682 transcripts), and protein coding genes (right, 54,181 transcripts). Each point represents the arithmetic mean across all experimental validated motifs for an RBP. **(C)** Cumulative distribution of RNA transcripts captured by each human RBP based on the position weight matrix for its motif, non-coding, and protein coding transcripts. **(D)** Amino acid alignment of RRM-containing RBPs to RRM1 of human PABPC4.

**Figure S12.**
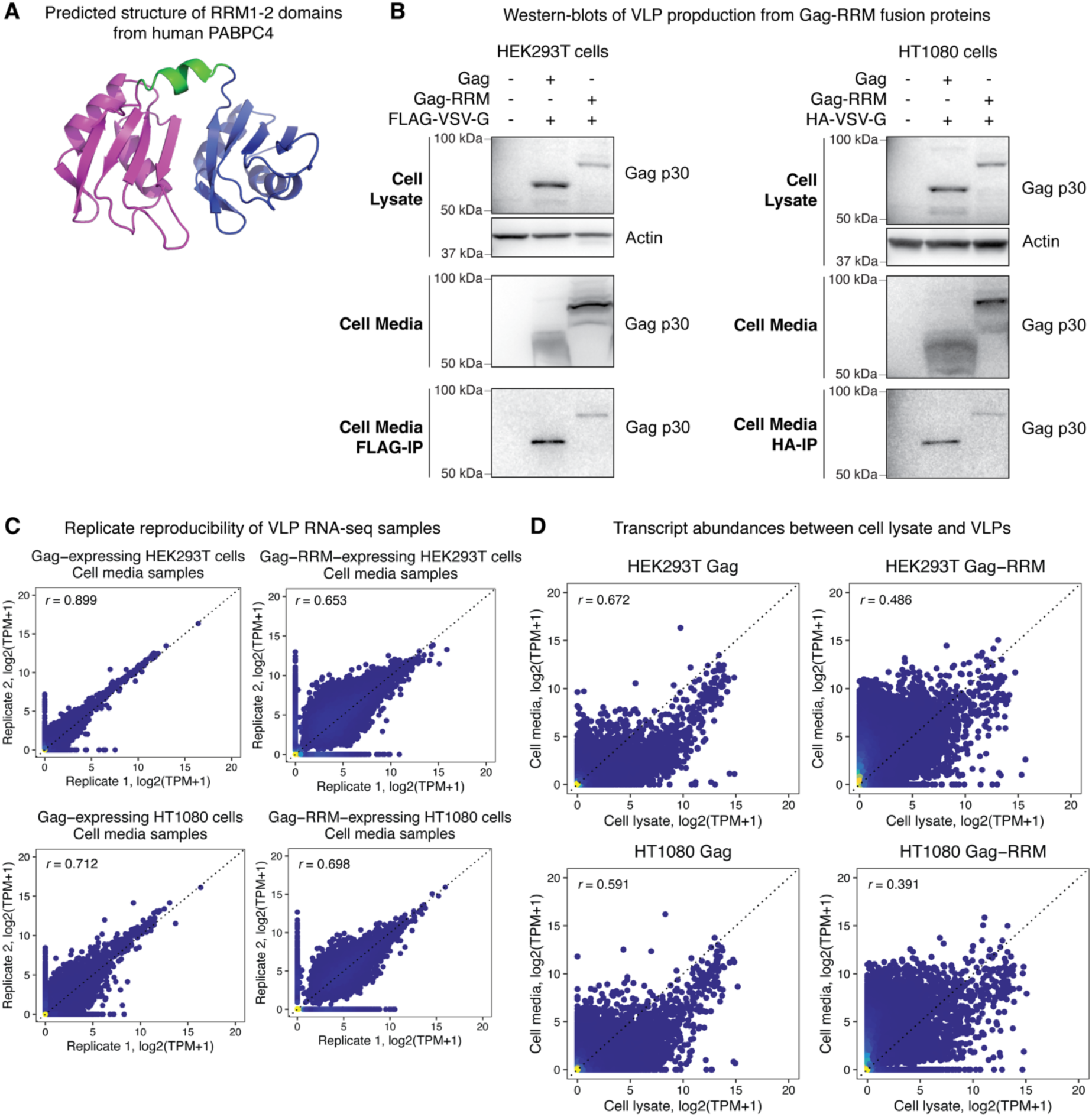
Characterization of self-reported RNA from Gag-RRM VLPs. **(A)** AlphaFold2-predicted structure of RRM1-2 from human PABPC4. **(B)** Western-blot verification that Gag-RRM fusion proteins form VLPs from HEK293T cells (left) and HT1080 cells (right). Cells were engineered to constitutively express FLAG-VSV-G and either Gag or Gag-RRM via lentiviral transduction. VLPs were isolated from the cell media by FLAG-IP following 48 hours of VLP production. Protein from cells and the culture media was immunoblotted for MLV Gag p30 in both cell lysate and media fractions, as well as for actin in cell lysate as a loading control. Expected molecular weights: 62.07 kDa for Gag. **(C)** Replicate reproducibility of transcript abundances from VLP RNA-seq samples of HEK293T and HT1080 cells stably expressing Gag or Gag-RRM. **(D)** Transcript abundances between cell lysate and VLPs samples from HEK293T and HT1080 cells expressing Gag or Gag-RRM. Pearson correlation coefficients are noted within each plot. TPM = transcripts per million.

**Figure S13.**
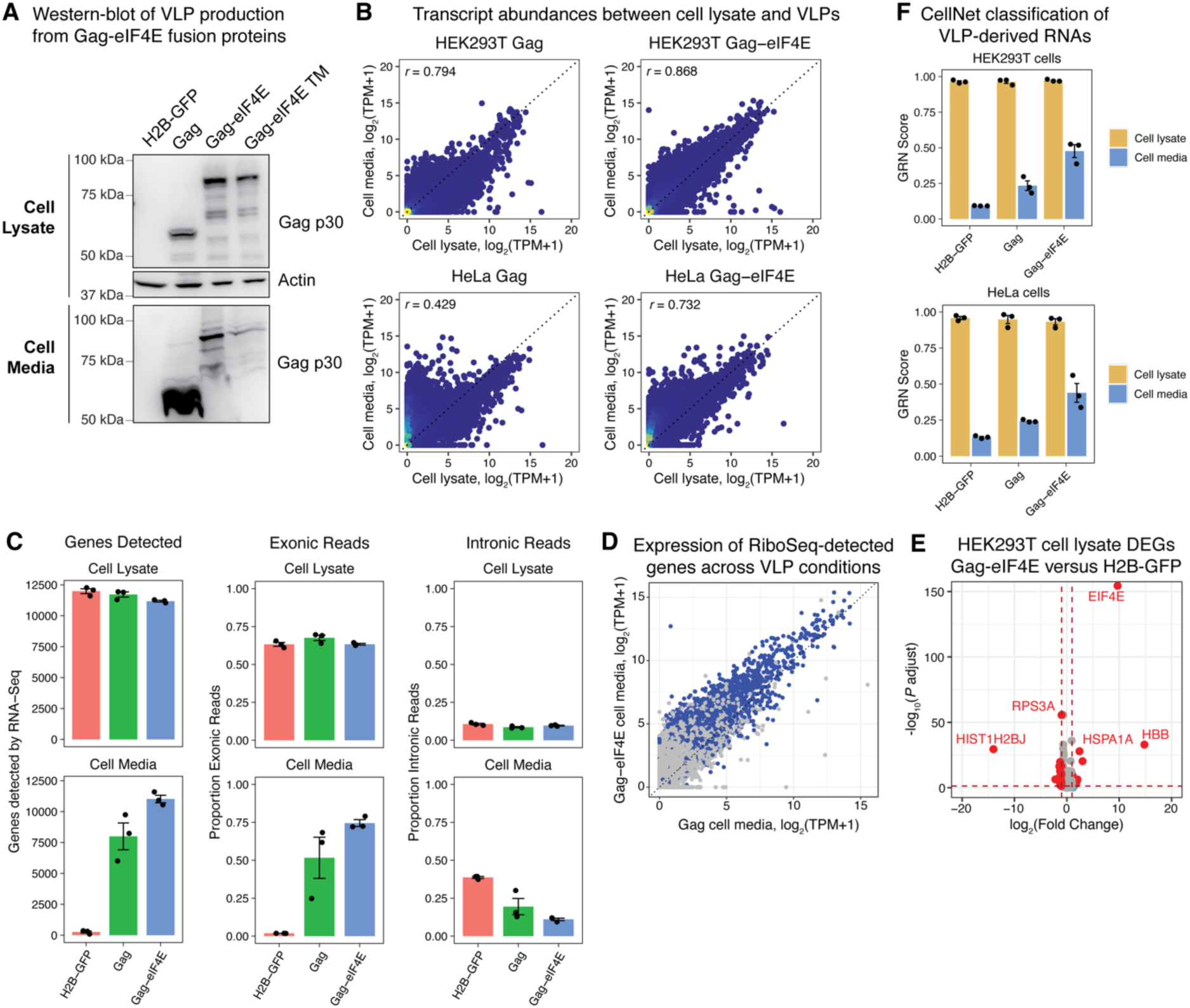
Characterization of self-reported RNAs from Gag-eIF4E VLPs. **(A)** Western-blot verification that Gag-eIF4E fusion proteins form VLPs. HEK293T cells were transfected with vectors to express either H2B-GFP, Gag, Gag-eIF4E, or Gag-eIF4E containing W56L, W102L and W166L mutations to disrupt 5′ cap binding (termed Gag-eIF4E TM). Protein from cells and the culture media was immunoblotted for MLV Gag p30 in both cell lysate and media fractions, as well as for actin in cell lysate as a loading control. Expected molecular weights: 62.07 kDa for Gag. Expected molecular weights: 62.07 kDa for Gag, 87.87 kDa for Gag-eIF4E, and 87.65 kDa for Gag-eIF4E TM. **(B)** Scatterplot of RNA transcript abundances from cellular lysate and Gag or Gag-eIF4E VLPs from HEK293T cells (top row) and HeLa cells (bottom row). Each point represents a gene detected from RNA-seq and the values are averaged over n=3 biological replicates. Pearson correlation coefficients between cellular lysate and VLPs are noted in each plot. **(C)** Quantification of the number of genes detected (left), proportion of exonic reads (middle), and proportion of intronic reads (right) from RNA-seq samples in **(B)**. **(D)** Scatterplot of transcript abundances of genes detected from Gag versus Gag-eIF4E VLPs produced from HEK293T cells. Genes undergoing active translation, as determined by Ribo-Seq from Clamer, *et al* 2018 (PMID: 30355487)^63^ are highlighted. **(E)** Volcano plot of DEGs in the cell lysates of Gag-eIF4E versus H2B-GFP expressing cells. Significant DEGs were defined as log2(fold change) < −1 or log2(fold change) > 1 and FDR-adjusted *P* value < 0.05. **(F)** CellNet GRN classification of cellular lysate and VLP-derived RNAs from cells expressing either H2B-GFP, Gag, or Gag-eIF4E. GRN scores from HEK293T cells (top) and HeLa cells (bottom) reflect the degree to which the samples recapitulate the HEK293T or HeLa GRNs in the CellNet classifier, respectively. n=3 replicates per condition. TPM = transcripts per million.

**Figure S14.**
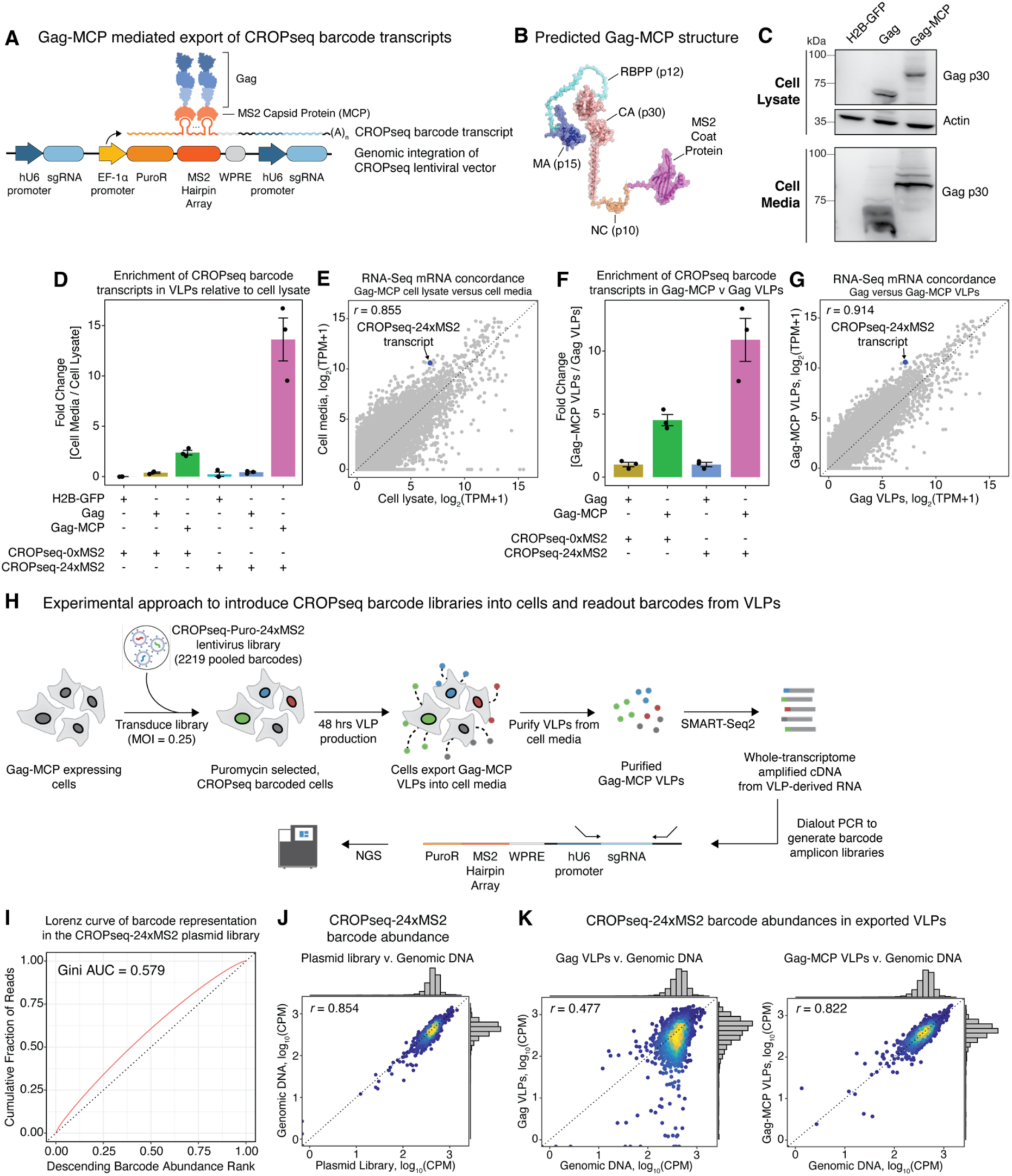
Gag-MCP mediated export of RNA barcode transcripts within VLPs. **(A)** Experimental schematic for directed export of RNA barcode transcripts within VLPs via MS2 containing CROPseq barcode transcripts and an engineered Gag fusion protein to the MS2 Coat Protein (MCP). The CROPseq vector is modified with 24x tandem repeats of MS2 hairpins within the RNA Pol II transcript driven by the EF1α promoter, which encodes the RNA barcode and is polyadenylated. The MS2 hairpins within the RNA barcode transcript can be bound by MCP fused to the C-terminus of Gag. Gag-MCP fusion proteins form into VLPs that package the RNA barcode transcript. **(B)** AlphaFold2-predicted structure of engineered Gag-MCP protein, whereby MCP (N55K variant) was fused to the C-terminus of MLV Gag via a 3x(G4S) linker. **(C)** Western-blot of HEK293T cells transfected with H2B-GFP, Gag, or Gag-MCP expression vectors. The cellular lysate and media were assayed for actin (loading control) and MLV Gag p30 antigen to determine production of VLPs. Expected molecular weights: 62.07 kDa for Gag and 76.87 kDa for Gag-MCP. **(D)** Enrichment of the CROPseq RNA barcode transcript in VLPs compared to cellular lysate. HEK293T cells lentivirally transduced with either CROPseq-0xMS2 or CROPseq-24xMS2 barcode vectors were transfected with H2B-GFP, Gag, or Gag-MCP expression vectors. RNA-seq libraries were prepared from cell lysate and VLPs purified from cell media collected 48 hours post-transfection. n=3 transfection replicates per condition. **(E)** Scatterplot of gene expression detected from Gag-MCP VLPs and the corresponding cell lysate. Each point represents a gene detected from RNA-seq and the values are averaged over n=3 matched replicates from cellular lysate and VLPs. The CROPseq RNA barcode transcript is highlighted and Pearson correlation of gene expression from cell lysate and Gag-MCP is noted as an inset. TPM = transcripts per million. **(F)** Same as **(D)** but CROPseq RNA barcode transcript expression enrichment is determined between Gag-MCP and Gag VLPs. **(G)** Same as **(E)** but with compared gene expression detected from Gag-MCP and Gag VLPs. **(H)** Experimental approach to introduce CROPseq barcode libraries into cells and readout barcodes from VLPs. **(I)** Lorenz curve of barcode representation in the CROPseq-24xMS2 plasmid library. **(J)** Barcode abundance quantified by NGS between the plasmid library and from genomic DNA following puromycin selection. **(K)** Barcode abundance between genomic DNA and Gag VLPs (left) or Gag-MCP (right). Pearson correlations are presented within the plots. CPM = counts per million.

## SUPPLEMENTAL TABLES

**Table S1.** Amino acid sequences of self-reporting constructs

**Table S2A:** Differentially expressed genes during iNGN differentiation detected from cell lysate: Day 4 versus Day 0

**Table S2B:** Differentially expressed genes during iNGN differentiation detected from VLPs: Day 4 versus Day 0

**Table S3.** List of human RBPs used for transcriptome-wide motif analyses

**Table S4.** Curated set of RBP motifs utilized for transcriptome-wide motif analysis and associated metadata

**Table S5.** Motif frequency quantification of RBPs in the human transcriptome

**Table S6.** Counts matrix of CROPseq barcodes from plasmid, genomic DNA and Gag-MCP VLP libraries

## RESOURCE AVAILABILITY

### Lead contact

Further information and requests for resources and reagents should be directed to the lead contact, Paul C. Blainey.

### Materials availability

Plasmids generated in this study are available from Addgene. All unique/stable reagents generated in this study are available from the lead contact with a completed Materials Transfer Agreement.

### Data and code availability

Any information required to reanalyze the data reported in this paper is available from the lead contact upon request

## METHOD DETAILS

### Cell culture

HEK293T (ThermoFisher Scientific cat. no. R70007), HT1080 (ATCC cat. no. CCL-121) and HeLa cells (a gift from Iain Cheeseman at the Whitehead Institute) were cultured in a humidified incubator at 5% CO_2_ at 37°C using high-glucose DMEM (Invitrogen cat. no. 10566016) supplemented with 10% FBS and 1% penicillin/streptomycin. Stromal fibroblasts were isolated from proliferative phase eutopic endometrial biopsies and all participants provided informed consent in accordance with a protocol approved by the Partners Human Research Committee and the Massachusetts Institute of Technology Committee on the Use of Humans as Experimental Subjects. Primary isolated endometrial stromal fibroblasts were routinely cultured at 37°C and 5% CO_2_ in phenol red-free DMEM/F12 media supplemented with 1% penicillin/streptomycin and 10% v/v dextran/charcoal treated fetal bovine serum (Atlanta Biologicals). Stromal fibroblasts were split 1:2 once they reached 70-80% confluency, and the medium was replaced every 2-3 days. Human iPS cells were cultured on plasticware coated with growth factor-reduced matrigel (Corning cat. no. 354230), diluted in DMEM-F/12 (Stem Cell Technologies cat. no. 36254) according to manufacturer instructions. The 1157.2 human iPS cell line (Boston Children’s Hospital Stem Cell Core) was maintained in StemFlex media (Stem Cell Technologies cat. no. A3349401), whereas the PGP1 iNGN iPS cell line (a gift from George Church at Harvard Medical School) was cultured in mTeSR medium (Stem Cell Technologies cat. no. 85850). iPS cells were propagated as colonies using ReleSR (Stem Cell Technologies cat. no. 100-0483) upon reaching 70% confluency. Cell cultures were regularly screened for mycoplasma contamination.

### Molecular cloning and construction of self-reporting plasmids

All plasmid vectors were designed in Benchling and generated by Gibson Assembly using NEB’s Gibson Assembly Master Mix, according to the manufacturer’s protocol. The MLV Gag ORF (UniProt ID: P03332) was synthesized by IDT as a gBlock and cloned in frame with a P2A-GFP sequence into the pcDNA3 vector in order to generate the episomal expression vector, pGag. As a negative control for VLP formation, we also created a Gag truncation mutant where the MA domain was deleted in order to abolish VLP maturation. We PCR dialed-out amino acids 216-530 from the MLV Gag ORF gBlock and equivalently cloned it in frame with a P2A-GFP sequence into the pcDNA3 vector in order to generate plasmid, pGagΔMA. We fused mCherry to the C-terminus of Gag via a 3x glycine-serine linker and cloned the ORF into pcDNA3 to create plasmid, pGag-mCherry. We utilized the pLenti vector to create all lentiviral expression plasmids. We PCR dialed out the Gag-P2A-GFP ORF from pGag and cloned it in frame with P2A-ZeoR into pLenti, downstream of a CAG promoter in order to create the tricistronic vector, pLV_CAG_Gag. We performed the comparable PCR dial out from pGagΔMA in order to create pLV_CAG_GagΔMA. Lentiviral vectors expressing Gag fusion proteins were generated by cloning ORFs encoding RNA binding proteins into pLV_CAG_Gag at the C-terminus of Gag with a 5x glycine-serine linker. The RRM1-2 domains from human PABPC4 (UniProt: Q6IQ30) was synthesized as a gBlock by IDT in order to create pLV_CAG_Gag-RRM. The eIF4E ORF was PCR dialed out from a whole-transcriptome amplification library in order to create pLV_CAG_Gag-EIF4E. We created a triple mutant variant of eIF4E by designing Gibson Assembly primers to substitute the tryptophans 56, 102 and 166 for leucines then PCR amplified the eIF4E cDNA ORF with said primers and assembled the fragments via Gibson Assembly to generate pLV_CAG_Gag-EIF4E-TM. Epitope-tagged VSV-G constructs were generated utilizing the VSV-G ORF from pMD2.G (a gift from D. Trono) and inserting a FLAG (DYKDDDDK) or HA (YPYDVPDYA) sequence following the signal peptide at amino acid position 23. FLAG or HA tags were inserted into tNGFR after the signal peptide and the ORFs were synthesized as dsDNA by Twist Biosciences. The epitope-tagged ORFs were cloned into pcDNA3 or pLenti backbones with a 2A-mCherry-2A-PuroR cassette to derive episomal or lentivirus vectors, respectively. Finally, piggyBac vectors for expression in iPS cells were generated with a piggyBac transposon backbone (System Biosciences) containing a CBX3 ubiquitous chromatin opening element (UCOE) upstream of an EF1ɑ promoter driving expression of Gag or GagΔMA in frame with a P2A ribosomal skipping peptide and mNeonGreen reporter.

All Gibson Assembly reactions were purified with a DNA Clean & Concentrator column (Zymo cat. no. D4033) prior to transformation into NEB Stable Competent *E. coli* (cat. no. C3040H). Transformed *E. coli* were grown overnight at 30°C on agar plates with 50 µg/mL carbenicillin selection. Individual colonies were picked for liquid culture in LB media supplemented with 100 µg/mL ampicillin and plasmid DNA was subsequently isolated using a QIAGEN Plasmid Plus Midi Kit with endotoxin removal. Plasmid sequences were fully verified by whole plasmid sequencing (GeneWiz or Primordium Labs, Inc). All plasmids have been deposited with Addgene.

### Transient transfection of self-reporting constructs

Cells were seeded into tissue culture plates at a density of 100,000 cells/cm^2^. The next day, cells were transfected with 3000 ng of total DNA comprising a Gag expression plasmid, VSV-G expression plasmid and pUC19 plasmid (2:3:4 ratio by mass) using Lipofectamine 3000 (Thermo Fisher). The media was changed 6 hours post-transfection and was collected for downstream processing 48 hours post-transfection.

### VLP purification from cell media

VLP-containing cell media from transient transfection experiments or stable Gag-expressing cells was first processed to remove cellular debris by centrifugation at 1,000 g for 5 minutes at 4°C. Next, the media was further filtered through a 0.45 μm cellulose acetate filter (VWR cat. no. 28145-481) and subsequently concentrated by centrifugation with a 100 kDa Amicon cutoff filter (Millipore Sigma cat. no. UFC5100) at 2500 g for 15 minutes at 4°C. The retentate within the filter was then either frozen at −80°C for storage or used immediately for subsequent assays.

For FLAG-based immunoprecipitation of VLPs, 20 µL of Anti-FLAG M2 Magnetic Beads (Sigma cat. no. M8823) were used per sample and washed 3x with TBS buffer. The beads were resuspended in 500 µL TBS + 1% Tween-20 and incubated with the VLP retentate on a rotisserie at 4°C overnight. The next day, the beads were washed 3x with TBS + 1% Tween-20 and eluted in 500 ng/uL 3x FLAG peptide (Sigma cat. no. F4799) at 1200 RPM shaking for 30 minutes at 4°C. For HA-based immunoprecipitation of VLPs, 50 µL of His-Tag Dynabeads (Invitrogen cat. no. 10103D) were used per sample according to the manufacturer’s protocol. HA-tagged VLPs were eluted in 50 µL of elution buffer (300 mM imidazole, 50 mM sodium phosphate, 300 mM NaCl and 0.01% Tween-20) at 1200 RPM shaking for 30 minutes at 4°C. The eluted VLPs were used directly in downstream assays or stored at −80°C.

### Microscopy for intracellular localization of MLV Gag proteins

HEK293T cells were seeded into a 24-well glass-bottom culture plate at a density of 100,000 cells/cm^2^. The following morning, each well was transfected with 165 ng of plasmid encoding a Gag-mCherry fusion protein and 330 ng of pUC19 using Lipofectamine 3000 (Thermo Fisher), per the manufacturer’s instructions. 48 hours following the start of the transfection, the cells were fixed with 4% PFA for 15 minutes, permeabilized with 0.2% Triton-X for 20 minutes, and blocked with 1% BSA for 30 minutes at room temperature. The samples were then stained with mouse anti-EEA1 antibody (BD Biosciences, 1:500 dilution, RRID: AB_397830) in 1% BSA for 2 hours at room temperature. The cells were then incubated with goat anti-mouse IgG Alexa Fluor 647 (Thermo Fisher, 1:500 dilution, RRID: AB_2535804) and Alexa Fluor 488 phalloidin (Thermo Fisher, 1:500 dilution) in 1% BSA for 30 minutes at room temperature. The cells were then stained with 100 ng/mL DAPI in 2X SSC and imaged at 40X magnification.

### Scanning electron microscopy

VLPs were produced from HEK293T cells via plasmid transfection of MLV Gag expression plasmids, as described in “Transient transfection of self-reporting constructs.” After 48 hours-post transfection, the cell media was centrifuged at 2000 g for 15 minutes to remove debris followed by filtration through 0.45 μm cellulose-acetate filters (VWR cat. no. 28145-481). The cleared media was then ultracentrifuged at 100,000 g for 2 hours using a 20% sucrose cushion. Carbon-coated copper mesh grids were coated in 1 μg/mL Alcian Blue solution. Grids were washed with ultrapure water, dried and then incubated at room temperature with VLP-containing supernatants. SEM micrographs were acquired on a Zeiss Crossbeam 540 scanning electron microscope.

### RNA temporal stability in VLPs

We assessed the temporal stability of RNA within VLPs by first producing VLPs via transient transfection of MLV Gag into HEK293T cells with n=3 transfection replicates, as described above. VLPs were purified from cellular media by clearing cellular debris via centrifugation at 1000 g for 10 minutes at 4°C and then filtered through 0.45 µm cellulose acetate filters. The purified VLPs were then incubated at 37°C for 4 days and sampled every 24 hours in order to quantify GAPDH expression via RT-qPCR. RNA was extracted from VLPs using a Qiagen QIAamp Viral RNA Mini Kit and then treated with DNase I (NEB cat. no. M0303S) for 10 minutes at 37°C. The DNase I reaction was inactivated with 5 mM EDTA and then subsequently cleaned up with 1X RNA SPRI beads (Beckman Coulter cat. no. A63987). 500 ng of RNA per sample was reverse transcribed using an iScript cDNA Synthesis kit (Bio-Rad cat. no. 1708890). GAPDH expression was quantified with a sensitive TaqMan-based qPCR assay using a commercial primer/probe set from ThermoFisher Scientific (Hs02758991_g1). Briefly, we first made a PCR standard by gel extracting the 93-bp PCR amplicon produced from the GAPDH primer/probe set using a 2% agarose gel and Zymo DNA Gel Extraction kit (Zymo cat. no. D4007). The concentration of gel-extracted GAPDH amplicon was quantified with a QuBit high sensitivity dsDNA assay and then diluted in nuclease-free water to create 10-fold serial dilutions ranging from 1e9 to 1e2 molecules/uL. Standards, cDNA samples and no-template controls were assayed by RT-qPCR in technical triplicate using JumpStart Taq DNA polymerase ReadyMix (Sigma cat. no. P2893), with Rox as a background control dye and the following thermal program: 94°C for 2 minutes, and 35 cycles of 94°C for 30 seconds, 60°C for 30 seconds and 72°C for 30 seconds. The PCR standard curve exhibited high technical reproducibility and a coefficient of determination of R^2^ = 0.9962. GAPDH copy number expression for each technical replicate was determined from the PCR standard curve and expression for each sample is the arithmetic mean across all technical replicates. To determine temporal stability, the copy number expression of GAPDH at each time-point was normalized to the starting Day 0 copy number baseline expression.

### Nuclease treatment of VLPs

HEK293T cells were transiently transfected in 6-well plates (as described above) with plasmids to express H2B-GFP, GagΔMA or Gag, and iPS cells were engineered via piggyBac transposition to stably express GFP, GagΔMA or Gag. Cells were incubated for 48 hours to allow for VLP secretion into the cell media. The media was then collected from cells and treated with 200 U/mL benzonase endonuclease (Sigma cat. no. E1014) at 37°C for 60 minutes with mixing every 15 minutes. Cellular debris was then cleared by centrifugation at 1,000 g for 5 minutes at 4°C and then further filtered through a 0.45 μm cellulose acetate filter (VWR cat. no. 28145-481). The cleared media was then concentrated with 100 kDa Amicon cutoff filter (Millipore Sigma cat. no. UFC5100) at 2500 g for 15 minutes at 4°C. The retentate was then lysed with a 2X TCL lysis buffer (Qiagen cat. no. 1070498) or treated with TRIzol to isolate RNA and then used for downstream qPCR assays or RNA-seq library construction.

### Protein structure predictions

We utilized ColabFold, a Google cloud implementation of AlphaFold2^65,66^ to predict the structure of engineered variants of MLV Gag and epitope-tagged envelope proteins. For all predictions, amino acid sequences were queried as monomers in AlphaFold2_mmseqs2 (v1.5.5) with default model/MSA settings and num_recycles set to 6. To model the interaction of the 5′ mRNA cap with either wild-type eIF4E or eIF4E exhibiting W56L, W102L, W166L mutations, we utilized AlphaFold3^62^, implemented through AlphaFold Server (alphafoldserver.com), with the amino acid sequence of the proteins and guanosine-5′-diphosphate as a ligand. The resulting PDB file for each predicted protein structure was visualized with PyMOL (v2.5.7).

### Lentivirus production

HEK293FT cells were seeded at 1e6 cells/well in 6-well plates. The following day, cells were transfected when 90-95% confluent with pMD2.G (Addgene #12259), psPAX2 (Addgene #12260), and a lentiviral transfer plasmid (2:3:4 ratio by mass, 3 µg total plasmid) using Lipofectamine 3000 (Thermo Scientific cat. no. L3000015). Media was exchanged after 6 hours and viral supernatant was harvested 48 hours after transfection and filtered through 0.45 μm cellulose-acetate filters (VWR cat. no. 28145-481). The purified lentivirus was frozen at −80°C until needed for infection.

### Generation of constitutive MLV Gag expressing cells

Constitutively self-reporting HEK293T cells, HT1080 cells and primary stromal fibroblasts were generated by lentiviral transduction of MLV *gag*. For some experiments, cells were then subsequently infected with lentivirus to introduce epitope-tagged envelope proteins. Lentiviruses were produced using plasmids pLV_CAG_GagΔMA or pLV_CAG_Gag (as described above). Cells were then transduced with lentiviruses in the presence of 8 µg/mL polybrene in order to achieve ∼30% fluorescent reporter-positive cells for single-copy integrations. The cells were incubated in virus for 24 hours followed by antibiotic selection 48 hours post-transduction at the following concentrations: 1 μg/mL puromycin (Thermo Scientific A1113802), and 200 μg/mL zeocin (Thermo Scientific R25001). Cells were under antibiotic selection until uninfected control cells died (∼5 days for puromycin and ∼9 days for zeocin).

Stable Gag-expressing iPS cells were generated through piggyBac transposition to achieve long-term transgene expression and avoid transgene silencing from lentiviral vectors. iPS cells were single cell dissociated with TrypLE reagent (ThermoFisher Scientific cat. no. 12604013) and 800,000 cells were nucleofected with 2.5 µg of Super piggyBac Transposase (SBI cat. no. PB200A-1) and 10 µg of transposon plasmid using Lonza Cell Line Nucleofector Kit V (Lonza cat. no. VCA-1003) on an Amaxa IIb Nucleofector with program B-016. iPS cells post-nucleofection were cultured with ROCK inhibitor, RevitaCell (ThermoFisher Scientific cat. no. A2644501) for 24 hours and FACS sorted one week later to enrich for mNeon+ cells.

### CellTiter-Glo time-course assay

We utilized CellTiter-Glo assays to determine cell growth kinetics from constitutively self-reporting cells. First, 5,000 cells were seeded per well in Nuncleon white polystyrene 96-well assay plates (Thermo Scientific cat. no. 136101), avoiding edge wells. Each condition was prepared in technical triplicate and replicate plates were prepared at the start of the experiment for each timepoint in the time-course. The CellTiter-Glo assay (Promega cat. no. G7572) was performed according to the manufacturer’s protocol on a BioTek Synergy Neo microplate reader. The raw luminescence signal for each timepoint was normalized to the starting Day 0 baseline signal.

### RNA-Sequencing

RNA from cells, cell media, or IP-purified VLPs was extracted using 2X TCL lysis buffer (Qiagen cat. no. 1070498). At least two technical replicates were prepared per sample using the SMART-Seq2 protocol as previously published^46^ with some modifications. Briefly, the lysed samples were cleaned with 2.2X RNA SPRI beads (Beckman Coulter cat. no. A63987) and reverse transcribed in the presence of a template switching oligo (Exiqon) with Maxima RNase H-minus RT (Thermo Fisher Scientific cat. no. EP0751) using a polyT primer containing the ISPCR sequence. Whole transcriptome amplification proceeded with KAPA HiFi HotStart ReadyMix using an ISPCR primer according to the following thermal program: 98°C for 3 minutes, 27 cycles of 98°C for 15 seconds, 67°C for 20 seconds, and 72°C for 6 minutes, and a final extension step of 72°C for 5 minutes. The amplified cDNA was cleaned with 0.8X DNA SPRI beads (Beckman Coulter cat. no. B23318). Ten nanograms of DNA was tagmented at 58°C for 10 minutes in a 10 µL reaction containing 2 µL of 5X tagmentation buffer (50 mM Tris-HCl, 25 mM MgCl2 pH 8.0), 2 µL of Tris Buffer (10 mM Tris-HCl, 1% Tween-20 pH 8), and 4 µL Nextera (Illumina). The reaction was stopped with 1% SDS and incubated at 72C for 10 minutes, then 4°C for 3 minutes. The tagmented library was cleaned with 1X DNA SPRI beads followed by 12 cycles of PCR with NEBNext High Fidelity polymerase to incorporate sample index barcodes and Illumina flow cell handles. The final libraries were pooled, diluted and sequenced on a NextSeq-500 (Illumina) in paired-end mode using a 75 cycle High Output Kit v2.

### RNA-Seq processing and analysis

RNA-seq paired-end reads were aligned to the hg19 reference transcriptome in order to quantify transcriptome-wide gene expression. We assessed the concordance of gene expression measurements with various NGS alignment tools, specifically STAR aligner (version 2.6.0c)^67^ and kallisto (version 0.46.0)^68^. RNA-seq reads were randomly downsampled prior to alignment to maintain equivalent sequencing depth across samples for analyses comparing different Gag VLP conditions. BAM alignment files were produced using STAR with command line options – outSAMtype BAM SortedByCoordinate –twopassMode Basic and –outSAMstrandField intronMotif. Read counts were calculated with featureCounts using uniquely mapped reads. Sample quality and alignment metrics were determined with RNA-SeQC^69^ using STAR-produced BAM files and an hg19 transcriptome GTF file retrieved from the UCSC Genome Browser. For Kallisto-based analyses, the hg19 cDNA fasta reference from the UCSC Genome Browser was appended with the coding sequences of Gag, GagΔMA, mNeon and VSV-G in order to create a custom Kallisto index via the “kallisto index” command. Transcript-level estimates were summed using the tximport package (version 1.28.0) in R (version 4.3.1). Differential expression was performed using the DESeq2 package (version 1.40.2)^70^ on the estimated counts matrix from Kallisto. Statistically significant genes varying between two conditions were identified using an FDR based *P* value adjustment method and alpha = 0.05. We defined genes as significantly differentially expressed if FDR < 0.05 and log2(fold-change) > 1 or log2(fold-change) < −1. Gene set enrichment analysis was performed using the clusterProfiler package (version 4.8.2) and unfiltered gene lists from DESeq2 ranked by log2(fold-change).

### Cell type classification and gene regulatory network inference with CellNet

We utilized CellNet^47^ to derive active gene regulatory networks (GRNs) from RNA-seq samples of cell lysate and VLPs, and to assess the ability of VLP-derived RNAs to reflect the GRNs in the lysate of the host cell. We retrained the CellNet classifier to incorporate GRNs reflective of HEK293T, HT1080, HeLa or primary stromal fibroblast cells. First, we quantified transcript abundances in cell lysate samples with cn_salmon() using a pre-prepared Salmon transcript index, salmon.index.human.122116.tgz available from https://github.com/pcahan1/CellNet. We constructed new cell-type specific GRNs with cn_make_grn() using samples from the June 20, 2017 edition of the human CellNet Processor (https://github.com/pcahan1/CellNet) and the cell lysate samples. We assessed the random forest classifier using cn_splitMakeAssess() and generated a new CellNet processor object with cn_make_processor(). We then applied CellNet with this retrained classifier using default settings to lysate and VLP RNA-seq samples and plotted the sample classification scores in R.

### Single cell export rate measurements

HEK293T cells were engineered with a dox-inducible Gag vector containing a GFP translation reporter via lentivirus infection. Cells were incubated with doxycycline (1 µg/mL) for 24 hours prior to FACS in order to induce production of VLPs. Single HEK293T cells in the top quartile of GFP expression were FACS sorted into a 384-well plate (Corning cat. no. 8794BC) containing 10 µL of dox-containing media per well using a Sony SH800Z cell sorter. No-dox control cells were single cell FACS sorted based on single events identified from the FSC-A and BSC-A event gate. The plate was centrifuged at 200 g for 1 minute at room temperature after sorting and cultured at 37°C. After 24 hours, brightfield images were acquired for each well in the culture plate using a Nikon Ti2 microscope to verify the presence of a viable cell. Cell media was then harvested from each well, lysed with 2xTCL buffer (Qiagen cat. no. 1070498) and imaged again to determine if cells were inadvertently aspirated as a result of cell media collection. The corresponding cell lysates were prepared by using 10 µL of 1xTCL (Qiagen cat. no. 1031576). RNA-seq libraries were generated using SMART-Seq2, as previously described. Cell lysate and media samples were pooled and paired-end sequenced if a cell was identified in both rounds of brightfield imaging during sample preparation. Paired-end sequencing reads were pseudo-aligned with kallisto (version 0.43.1) using a custom reference generated by concatenating the MLV Gag ORF, human cDNA reference transcriptome, and bovine cDNA reference transcriptome with an index build from 31-mers. For low-input and single-cell measurements from HT1080 cells constitutively expressing Gag, cells were similarly sorted, followed by cell lysate and cell media collection for RNA-seq as described above. Sequencing reads were aligned using STAR (version 2.6.0c) and read counts were calculated with featureCounts. Cell media samples were only considered for analysis if the corresponding cell lysates had at least 1,000 unique transcripts detected, indicating the cell was present in the lysate sample (not the supernatant sample). For conservative estimation of export rates, we assumed that each unique gene transcript detected was a unique RNA molecule measured.

### Immunoblotting for VLPs

The cell media from Gag-expressing cells was collected and processed as described in the section “VLP purification from cell media.” The cells were then washed once with PBS and incubated with TrypLE (Gibco) in order to create a single cell suspension. An aliquot of cells were resuspended in a protein extraction solution (150 mM NaCl, 1% Triton X-100, 50 mM Tris-HCl pH 7.5) supplemented with a protease inhibitor (Sigma cat. no. 4693159001) for 30 minutes at 4°C on a rotisserie. The cell lysate was centrifuged at 10,000 g for 5 minutes at 4C to pellet debris, then frozen at −80°C or used directly for immunoblotting. Cell protein lysate, purified cellular media, and/or IP-purified VLPs were reduced using 2X Tris-Glycine SDS Sample Buffer (Life Technologies cat. no. LC2676) and 10X NuPAGE Reducing Agent (Life Technologies cat. no. NP0009) at 98°C for 5 minutes. The reduced samples were then run on a Novex WedgeWell 8-16% Tris-Glycine Gel (Life Technologies cat. no. XP08165BOX) for 50 minutes at 225V. The gel was then transferred to a PVDF membrane (Life Technologies cat. no. IB24002) using an iBlot 2 device (Life Technologies). The PVDF membrane was blocked in 5% milk (Bio-Rad cat. no. 1706404XTU) for 1 hour at room temperature with gentle shaking, followed by an overnight incubation at 4°C with gentle shaking in primary antibodies against Gag at 1:2000 dilution (Abcam cat. no. ab100970, RRID: AB_10696216) and Actin at 1:5000 dilution (Abcam cat. no. ab179467, RRID: AB_2737344). The next day the membrane was washed three times with 5% milk and incubated for 4 hours at 4°C in an anti-rabbit secondary antibody (Thermo Fisher Scientific, A32733, RRID: AB_2633282, 1:10,000 dilution). In experiments were we utilized GagΔMA-expressing cells as a VLP control, we stained for actin with a primary mouse antibody (Thermo Fisher Scientific, MA5-11869, RRID: AB_11004139, 1:400 dilution) and anti-mouse secondary (Thermo Fisher Scientific, A11001, RRID: AB_2534069, 1:10,000 dilution) to distinguish the similarly sized proteins The membranes were imaged on a Azure Biosystems C600 Imaging System and analyzed in Fiji (v2.14.0/1.54f).

### Live-cell iPS-to-neuron differentiation transcriptional time-course

To study the transcriptional dynamics of neurogenesis, we employed the PGP1 iNGN human iPS cell line (courtesy of Dr. George Church) expressing the transcription factors Neurogenin-1 and Neurogenin-2 under the control of a doxycycline-inducible promoter^48^. PGP1 iNGN cells were cultured on tissue culture plates coated with hESC-qualified Matrigel (Corning, cat. no. 354277) and grow in mTeSR media (StemCell Technologies, cat. no. 85850) with daily media changes. Cells were propagated as colonies with ReLeSR (StemCell Technologies, cat. no. 100-0483) upon reaching 70% confluency. We engineered PGP1 iNGN cells with self-reporting vectors via piggyBac transposition. Briefly, 1 million cells were singularized with TrypLE and nucleofected with 10 μg of transposon plasmid and 2.5 μg of transposase plasmid using Lonza Cell Line Nucleofection Kit V and program B-016 on an Amaxa IIb nucleofector. The cells were then cultured for 24 hours with RevitaCell (Thermo Fisher Scientific, cat. no. A2644501), after which fresh media was replenished daily as the cells recovered and expanded. mNeon+ cells were FACS sorted to isolate stably transposed cells, expanded, and then utilized for downstream experiments. The self-reporting PGP1 iNGN iPS cells were differentiated to bipolar-like neurons by first seeding 45 colonies/cm^2^ in a 15-cm dish on Day −1. We equivalently plated cells in multiple wells of a 24-well plate in order to destructively sample a well at each time-point of the differentiation for cell lysate control measurements. The following day (Day 0), the media was collected and purified for VLPs, the cells were washed with PBS to remove residual VLPs from the prior time-point, and then mTeSR media containing 0.5 μg/mL doxycycline (Sigma, cat. no. D9891-5G) was administered to initiate differentiation. Cells were destructively sampled for cell lysate control measurements by first washing the cells once with PBS, followed by singularization with TrypLE and lysis with TCL buffer (Qiagen). This process was repeated every 24 hours over 4 days of differentiation. Cell surface expression of NCAM1 was assayed via flow cytometry at the terminal time-point to determine the efficiency of the differentiation across Gag, GagΔMA and mNeon expressing PGP1 iNGN iPS cells (BioLegend cat. no. 362546, RRID: AB_2565964, 1:100 dilution, staining for 20 minutes at room temperature). Two independent differentiation experiments were performed for biological replication. VLPs were purified from cell media by first incubating the crude media with 100 U/mL benzonase at 37°C for 60 minutes with periodic mixing every 15 minutes. The media was centrifuged at 1000 g for 5 minutes, filtered through a 0.45 μm cellulose-acetate filter and then concentrated with a 100 kDa Amicon filter (Sigma, cat. no. UFC910008) at 2500 g for 20 minutes at 4°C. The retentate was lysed with 2xTCL buffer (Qiagen) and immediately frozen at −80°C. RNA-seq libraries were prepared with technical replication on cell lysate and VLP samples via SMART-Seq2 (as previously described above) and sequenced on an Illumina NextSeq in read configuration 75-8-8-75.

The hg19 UCSC reference transcriptome was appended to the coding sequences of mouse *Neurog1* and *Neurog2*, Gag, and mNeon to generate a custom kallisto index. RNA-seq paired-end reads were pseudoaligned to the custom index with kallisto (v.0.46.0). Differential gene expression analysis was performed using DESeq2 (version 1.44.0) with R (version 4.4.1) on the estimated count matrix output from kallisto, filtering for genes with at least 10 reads in 2 or more samples. Gene set enrichment analysis^71^ was performed using GSEApy on the log2(fold change) calculated from DESeq2. CellNet was applied to RNA-seq data from all time-points using the June 20, 2017 edition of the human CellNet Processor (available from https://github.com/pcahan1/CellNet) and without any further modifications. Embryonic stem cell GRN scores were then visualized as a function of time for cell lysate and VLP RNA-seq samples. All other downstream analysis was performed with custom Python scripts.

### *In silico* screening of RNA binding proteins

We sought to characterize the types and quantity of RNA transcripts that could be putatively bound by RNA binding proteins (RBPs) in the human genome, with the objective to nominate RBPs for fusion to Gag in order enhance RNA export in VLPs. First, we compiled a comprehensive, non-redundant list of human RBPs by aggregating publicly available databases^53–55^. We selected RBPs for inclusion if the RBP has an experimentally reported motif. We also annotated each RBP with molecular information from the RBP2GO^72^ and RBPWorld^73^ databases. We pulled the amino acid sequences for each RBP using the getUniProt() function from the protr package (version 1.7-2) in R and annotated if the amino acid sequence contains subcellular protein localization sequences. Position weight matrices (PWMs) of motifs for each RBP were downloaded from CisBP-RNA, ATtRACT and oRNAment databases. We converted the PWMs into a log2-likelihood representation with a pseudocount of 0.8 and compiled all motifs into a PWMatrixList using the TFBSTools package (version 1.38.0) in R. RBP motifs were selected for inclusion if the motif was experimentally determined using a non-mutated, wild-type form of the RBP. To determine RBP motif occurrences in the human transcriptome, we imported the hg19 reference transcriptome from the UCSC Genome Browser (https://hgdownload.soe.ucsc.edu/goldenPath/hg19/bigZips/refMrna.fa.gz) as a DNAStringSet using the Biostrings package (version 2.68.1) and used motifmatchr (version 1.22.0) “matchMotifs(genome = “hg19”, bg = “genome”, out = “scores”)” with our RBP motif PWMatrixList. The number of RNA transcripts containing an RBP motif and the number of motif occurrences per transcript were averaged across all motifs of an RBP (if an RBP has multiple motifs). This motif occurrence analysis was performed on different classes of RNAs, specifically, coding and non-coding transcripts. Based on these analyses we also performed amino acid based alignments of RNA binding domains using annotated RRM protein domains from our compiled RBP list. Alignments were performed using the pairwise2 function in the Biopython package.

### Export of CROPseq transcripts in VLPs

To facilitate export of specific RNA transcripts, we employed the MS2 hairpin system derived from the MS2 bacteriophage. First, the MS2 coat protein (MCP) (N55K variant) ORF was PCR amplified from MS2-P65-HSF1_GFP (Addgene, 61423)^74^ and inserted with a serine-glycine linker via Gibson Assembly into pLV_CAG_Gag at the C-terminus of MLV Gag in order to generate the plasmid pLV_CAG_Gag-MCP. We also modified the CROPseq-Puro vector (Addgene, 127458)^64^ by inserting a 24x tandem array of MS2 hairpins derived from phage-cmv-cfp-24xms2 (Addgene, 40651)^75^ via restriction cloning in between the puromycin resistance ORF and WPRE in order to generate the entry vector CROPseq-Puro-24xMS2.

To assess enrichment of MS2-tagged CROPseq transcripts within VLPs, we cloned a sham 20-nt barcode sequence into the CROPseq-Puro-24xMS2 and original CROPseq-Puro entry vectors via Golden Gate cloning. Each vector was stably integrated into HEK293T cells via lentiviral infection, followed by 5 days of puromycin selection. Gag-MCP and wild-type Gag expression plasmids were then transfected into each CROPseq expressing cell line, and VLPs were collected and purified after 48 hours (as previously described). RNA-seq libraries were prepared via SMART-Seq2 on cell lysate and purified VLPs (as previously described) and sequenced on an Illumina MiniSeq in a 76-8-8-76 read configuration. A custom kallisto index was generated by appending the expected RNA polII CROPseq transcript driven from the EF1ɑ promoter to the hg19 UCSC reference transcriptome. RNA-seq reads were pseudoaligned with kallisto (v.0.46.0) to the custom index and the fold change in CROPseq transcript expression between cell lysate and VLPs, and between Gag-MCP and Gag VLPs was calculated using the normalized transcripts-per-million expression matrix.

### Pooled CROPseq barcode cloning, lentivirus production, infection and readout from VLPs

We generated a 2,219 barcode library by subsampling the Broad Institute’s Dolcetto sgRNA library^76^. The barcode library was synthesized as an oligo array (Agilent), amplified with 12 rounds of PCR using KAPA HiFi DNA polymerase (Roche cat. no. 07958935001) and purified with a DNA Clean & Concentrator column (Zymo cat. no. D4033). The amplified barcode sequences were cloned into the CROPseq-Puro-24xMS2 entry vector at a 1:100 (vector:insert) molar ratio via Golden Gate Assembly with Esp3I (Thermo Fisher Scientific, cat. no. FD0454) and T7 ligase (Enzymatics cat. no. L6020L). The Golden Gate reaction was purified via DNA precipitation with isopropanol and 500 ng was transformed into Endura ElectroCompetent Cells (Lucigen, catalog no. 60242-2) following the manufacturer’s instructions. After transformation, Endura cells recovered for 1 hour at 37°C and then were grown for 16 hours at 30°C with 100 µg/mL ampicillin selection. The barcode plasmid library was isolated using a Plasmid Plus Midi Kit (Qiagen, catalog no. 12943) and sequenced for barcode representation on an Illumina MiniSeq (read configuration 75-8-8-0).

To produce lentivirus, 22 million HEK293FT cells were seeded into a 15-cm dish, and the following morning the cells were transfected with 12.1 µg pMD2.G (Addgene #12259), 18.15 µg psPAX2 (Addgene #12260) and 24.2 µg of the CROPseq-24xMS2 barcode plasmid library using Lipofectamine 3000 transfection reagent (Thermo Fisher Scientific, cat. no. L3000075) per the manufacturer’s instructions. Following 48 hours, the virus-containing cell media was collected, centrifuged at 1000g for 5 minutes at 4°C to clear cellular debris, filtered through 0.45 μm PVDF filters (Millipore cat. no. SLHVR04NL) and concentrated with a 100 kDa Amicon centrifugal filter (Millipore cat. no. UFC910024) at 2000g for 20 minutes at 4°C. Lentiviral particles were subsequently aliquoted and frozen at −80°C.

The CROPseq barcode library was introduced into HEK293T cells via reverse infection. First, HEK293T cells were singularized with TrypLE (Gibco cat. no.12604013) and 2 million cells were seeded into a T75 flask with the CROPseq lentivirus pool (MOI = 0.25), supplemented with 8 µg/mL polybrene (Milipore Sigma cat. no. TR-1003-G). The cells were incubated with the lentivirus for 24 hours, after which 1 µg/mL of puromycin (Thermo Scientific Fisher, A1113803) was administered for 5 days to select for lentivirus infected cells. The puromycin resistant cells were expanded and genomic DNA was extracted using a custom extraction buffer (1 mM CaCl2, 3 mM MgCl2, 1 mM EDTA, 10 mM Tris-HCl, 1% Triton X-100, 0.2 mg/ml Proteinase K) then subjected to the following thermal program: 65°C for 10 minutes followed by 95°C for 15 minutes. Barcode sequences were amplified from genomic DNA using Q5 High Fidelity DNA polymerase (NEB cat. no. M0541S) with 28 cycles of PCR and the following dialout primers: forward 5′-ACACGACGCTCTTCCGATCTTCTTGTGGAAAGGACGAAAC and reverse 5′-CTGGAGTTCAGACGTGTGCTCTTCCGATCAAGCACCGACTCGGTGCCAC. Barcode amplicon libraries were sequenced on an Illumina MiniSeq (read configuration 75-8-8-0).

We then assessed detection of CROPseq barcodes within VLPs. The CROPseq barcoded cell population was seeded into 6-well plates (1 million cells per well) and the following day was transfected with 2000 ng of expression vector encoding either H2B-GFP, wild-type Gag, or Gag-MCP. We performed n=3 replicate transfections per condition with Lipofectamine-3000, according to the manufacturer’s protocol. Following 48 hours post transfection, VLPs were collected and purified, as previously described. RNA was extracted from VLPs and cells using a 2xTCL buffer (Qiagen) and RNA-seq libraries were prepared via SMART-Seq2. We then PCR dial-out the CROPseq barcode transcript from the SMART-Seq2 whole-transcriptome amplification library using Q5 High Fidelity DNA polymerase with 28 cycles of PCR and the dialout primers listed above. Barcode amplicon libraries and RNA-seq libraries were sequenced on an Illumina MiniSeq (read configuration 76-8-8-76).

We quantified the abundance of barcodes from the plasmid library, genomic DNA and RNA-seq libraries of cellular lysate and VLPs using custom scripts written in R. In brief, we scanned fastq reads for the “CACCG” TSS motif of the human U6 promoter and identified exact matches to our white-list of barcodes. Raw sgRNA counts were normalized to the total read depth in each sample using the cpm() function in the edgeR package. The representation of barcodes within each sample was determined by calculating a Gini AUC using the ineq package in R.

